# Engineered bacterial siderophore production accelerates rock weathering for carbon removal

**DOI:** 10.1101/2025.04.08.647837

**Authors:** Neil C. Dalvie, Amogh P. Jalihal, Abigail Fitzgibbon, Jan-Tobias Böhnke, Mohammed Hijaz, Quincey A. Justman, Steven J. Davis, Pamela A. Silver, Michael Springer

## Abstract

Silicate mineral weathering (dissolution) is a scalable strategy for capture and storage of CO_2_ but is too slow for industrial deployment. Bacteria can accelerate mineral dissolution by secreting siderophores, molecules that solubilize iron released from the mineral that would otherwise passivate the mineral surface. Here, we investigated how to deploy siderophore-producing bacteria at scale to continuously enhance dissolution of the mineral olivine. We demonstrated that natural genetic regulation precludes continuous siderophore production in mineral bioreactors. To overcome this limitation, we engineered the marine bacterium *Alteromonas macleodii* to enhance its production of siderophores, conferring a 2.6-fold increase in the rate of olivine dissolution. Life Cycle Analysis indicated that renewable feedstocks and minimal replenishment of modified cells are critical to achieve net CO_2_ removal at scale. With these guidelines, we constructed pilot-scale continuous mineral bioreactors that use unprocessed seawater and a renewable acetate feedstock. In reactors with engineered cells, we directly measured removal of 0.50 g CO_2_ per day from the air through alkalinity generation, while a control reactor without cells did not generate alkalinity in net. These platforms and bioprocess principles will inform the design of large-scale, sustainable, unit operations for alkaline mineral processing.

## Introduction

Enhanced weathering of silicate rocks could enable removal and long-term storage of atmospheric CO_2_ at large scales.^1–3^ Mafic and ultramafic minerals such as olivine generate alkalinity upon dissolution. In marine and terrestrial mediums, this alkalinity is primarily comprised of stable bicarbonate ions HCO_3_^-^, derived from atmospheric CO_2_ (Fig. 1). In ongoing field trials in which mafic rocks were distributed into soils or coastal environments, the extent of CO_2_ removal has been challenging to measure due to slow rock weathering rates.^4,5^ Tank-based mineral processing would allow control over hydrology and mineral surface area, simplifying measurement of mineral dissolution and CO_2_ capture and storage. Existing unit operations to dissolve minerals leverage chemical or biological acids, however, precluding generation of alkalinity.^6^ A tank-based unit operation to accelerate the dissolution of silicate minerals in a low-cost environmental medium like seawater without reliance on acids could enable large-scale, measurable CO_2_ storage on the decadal timescale.

**Fig. 1.**
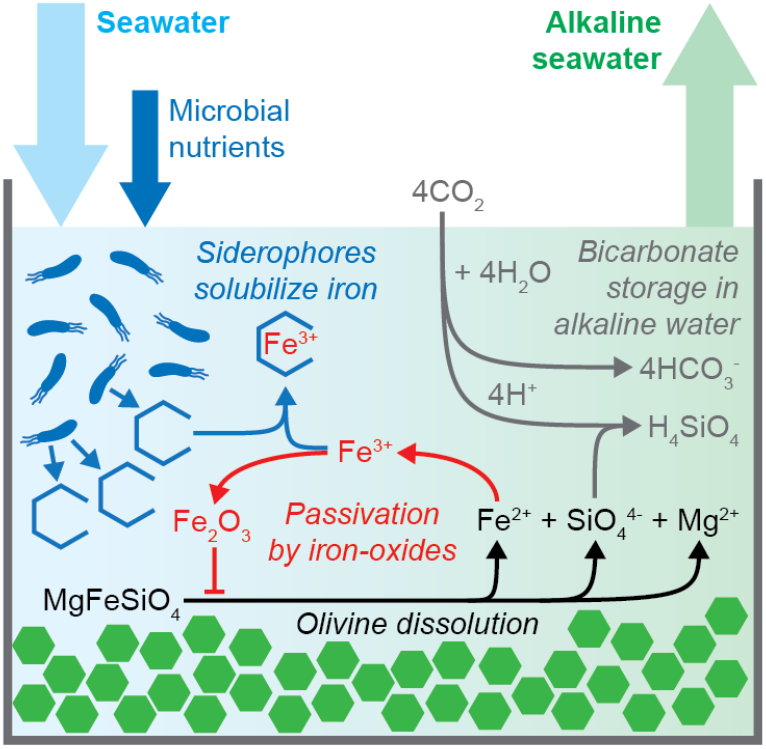
Bacterial siderophores accelerate olivine weathering. Siderophores solubilize ferric iron, preventing mineral passivation by iron oxides. After dissolution, silicate ions act as proton acceptors to stabilize carbonic acid in solution.

Bacterial siderophores, secreted secondary metabolites that chelate and solubilize ferric iron, can accelerate the dissolution of silicate minerals at neutral pH.^7^ Iron is abundant in mafic and ultramafic rocks and forms iron oxides in aqueous, oxidative conditions. During natural weathering, iron oxides precipitate back onto the mineral surface and prevent further mineral dissolution.^7^ Siderophores may prevent this passivation by solubilizing ferric iron. They have also been shown to directly extract iron from iron oxide minerals, a process that likely occurs in soils.^8,9^ In batch reactions, the siderophore desferrioxamine accelerated the dissolution of olivine by solubilization of passivating iron oxides.^10^ It was suggested, however, that siderophore production may be limited by the presence of olivine, preventing biologically accelerated mineral dissolution at large scales.

In this study, we developed continuous mineral bioreactors and experimentally confirmed that siderophore production is limited at scale. Then, we engineered a marine bacterium with enhanced siderophore production that can continuously accelerate olivine dissolution. We performed a Life Cycle Analysis (LCA) to assess the potential of mineral bioreactors to capture and store CO_2_ at industrial scale. Informed by our LCA, we constructed pilot-scale mineral bioreactors and demonstrated sustained acceleration of olivine dissolution by engineered bacteria in unprocessed seawater.

## Results

We sought to develop a scalable, tank-based unit operation for biologically accelerated olivine dissolution (Fig. 1). We reasoned that seawater was the optimal process medium; seawater is abundant, inexpensive, and can be returned to the environment carrying alkaline ions for long-term CO_2_ storage. Siderophore-producing bacteria are also ubiquitous in the ocean.^11^ Here, we used the marine bacterium *Alteromonas macleodii* which naturally produces the siderophore petrobactin.^12^ The genes for petrobactin synthesis were previously shown to be expressed in response to iron limitation,^13^ as is typical for most siderophore-producing bacteria (Fig. S1).^14,15^ In batch cultures with synthetic seawater medium, *A. macleodii* acquired iron from olivine for growth (Fig. S2). We could only achieve iron-limited growth, however, with 0.4 g/L of olivine. Ions released from this small quantity of olivine were not detectable in the seawater medium. To overcome this measurement challenge, we developed a platform for continuous cultures.

### Continuous mineral bioreactors

We developed parallelized bioreactors that enable continuous cell growth on a mineral substrate. In these modified eVOLVER chemostats,^16^ *A. macleodii* culture is continuously diluted while a fixed quantity of olivine sand is retained in the vessel via settling (Fig. 2A). After 1-2 days of continuous dilution, reactors with olivine sand reached steady state growth, while reactors without olivine exhibited washout (drop to near-zero cell density), supporting the assumption that cells were dependent on iron acquired from the olivine (Fig. S3).

**Fig. 2.**
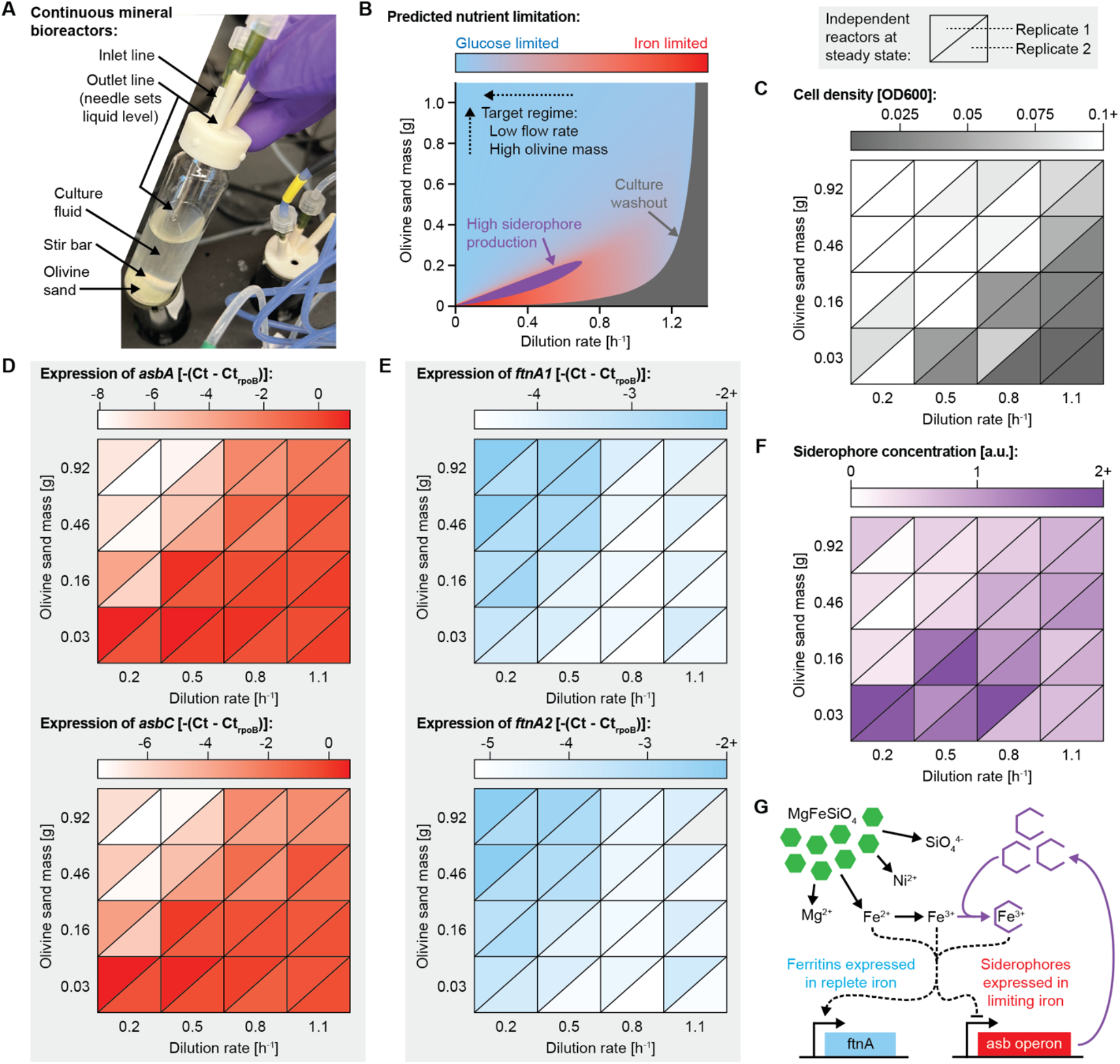
Olivine dissolution suppresses siderophore production. A) Continuous mineral bioreactors. Seawater media was added and removed at a constant rate while olivine sand remained in the vial by sinking. Stirring ensured well mixed culture fluid. B) Predictions for mineral reactors at steady state according to the theoretical chemostat model. Each point in this phase diagram represents a hypothetical reactor at steady state with a given dilution rate and mineral mass. Color spectrum represents the ratio of glucose to iron limitation according to the model. The purple region indicates process conditions that are predicted to yield a high steady state concentration of siderophores. C-F) Real reactors were operated with different dilution rates and mineral masses. The same set of 16 reactors were operated in replicate several weeks apart. Each triangle represents one chemostat at steady state. C) Optical density at 600 nm. D) Expression of siderophore synthesis genes *asbA* and *asbC*, measured by qPCR and normalized to a housekeeping gene *rpoB*. E) Expression of iron storage genes *ftnA1* and *ftnA2*. F) Concentration of the siderophore petrobactin measured by LCMS and normalized within each experiment. G) Iron released by olivine inhibits siderophore production.

Continuous mineral bioreactors also enable measurement of mineral dissolution at steady state. We developed a method to measure the total quantity of metal ions released from the olivine substrate by complete digestion of culture fluid in strong acid (Fig. S4A). Testing the method with synthetic metal mixtures revealed that Ni and Fe were good markers of olivine dissolution at small scale (Fig. S4B). Mg is a good marker of olivine dissolution at higher concentrations, but at low concentrations does not overcome the background signal of Mg in the seawater medium. In our pilot experiment, the rate of release of multiple metals from olivine did not change between two dilution rates or in the presence or absence of *A. macleodii* cells (Fig. S3). We hypothesized that cells were not iron-limited in these initial conditions and therefore did not produce siderophores.

We then sought to identify process conditions in which cells would experience continuous iron limitation and continuously produce siderophores. To guide our experiments, we modeled our continuous mineral bioreactors as chemostats (Appendix 1). In classical chemostats, cell growth is limited by nutrient influx (glucose in our system). Torres et al. previously modeled mineral chemostats, in which cell growth is limited instead by the release of iron from a solid mineral substrate.^10^ Our model builds upon this previous work by allowing cells to be limited by either glucose or iron, and by allowing control over both the mass of olivine in the vessel and the dilution rate of the medium. After solving our model, we predicted that cell growth at steady state would be primarily limited by iron only with low olivine mass and a high enough dilution rate (Fig. 2B, red). More specifically, we predicted that continuous siderophore production would only occur in a narrow range of process conditions in which cells experience iron limitation, but not so much that the culture washes out (Fig. 2B, purple).

We confirmed in real bioreactors that continuous iron limitation was limited to a narrow range of process conditions. We operated 16 parallel reactors with varied olivine masses and dilution rates in duplicate (Fig. S5). As predicted, cultures washed out above a certain dilution rate (Fig. 2C). In reactors with less olivine, washout occurred at lower dilution rates than in chemostats with more olivine, likely because less iron is available. We also measured the expression of genes that are markers for iron limitation.^13^ Siderophore synthesis genes were expressed at a high level with low olivine mass and high dilution rates, suggesting that growth was limited by iron in this regime (Fig. 2D). Conversely, iron storage genes were expressed at a high level with high olivine mass and low dilution rates, suggesting that growth was not limited by iron in this regime (Fig. 2E). We measured the relative quantity of the siderophore petrobactin and observed high concentrations only in conditions with iron limitation and high cell density (Fig. 2F). This observation corroborates our and others’ prediction that olivine could inhibit siderophore production at large scales (Fig. 2G).^10^ Indeed, we only observed increased release of Ni, Fe, and Si from olivine in chemostats with low olivine mass (Fig. S3). Together, these observations demonstrate that endogenous regulation in *A. macleodii* precludes siderophore production in bioreactors under reasonable operating parameters at industrial scale.

### Genetic engineering for siderophore production

To enable siderophore production in the presence of high quantities of olivine, we genetically engineered *A. macleodii*. To enable this engineering, we characterized a set of synthetic, constitutive promoters (https://parts.igem.org/Promoters/Catalog/Anderson) and observed a >30-fold range of GFP fluorescence (Fig. S6). We then cloned the entire petrobactin synthesis operon *asb* into an episomal plasmid under control of each synthetic promoter and evaluated each transformed strain for petrobactin secretion by CAS agar assay (Fig. 3A).^17^ While most plasmids conferred little to no increase in CAS assay signal, the strain carrying a plasmid with the *asb* operon under control of the second strongest promoter (J23100) exhibited a dramatic increase in CAS assay signal, suggesting increased siderophore secretion and cellular iron uptake (Fig. 3B, Fig. S7). We denoted this strain “asb+P”. Interestingly, we were unable to successfully transform the plasmid with the *asb* operon under control of the strongest promoter (J23101). We hypothesize, therefore, that the asb+P strain is near the maximal expression level of the *asb* operon that would confer toxicity.

**Fig. 3.**
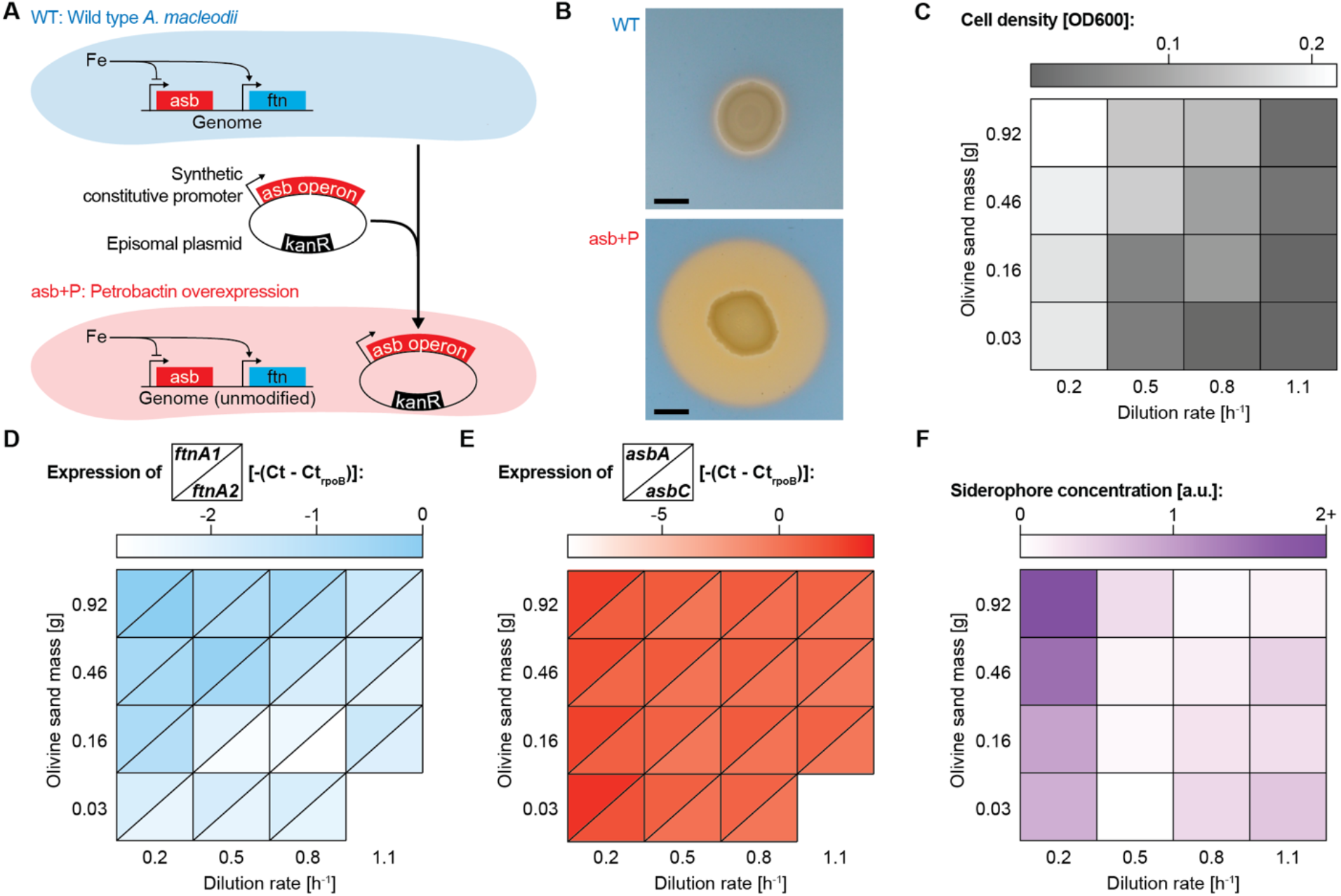
Engineered bacteria constitutively produce siderophores. A) Construction of the asb+P strain. The *asb* operon was cloned onto an episomal plasmid and conjugated back into *A. macleodii* under control of a synthetic constitutive promoter. B) Photo of cells growing on CAS agar plates. Yellow signal indicates sequestration of iron from the blue dye by secreted siderophores. Scale bar = 1 cm. C) Optical density of continuous mineral bioreactors at steady state with different dilution rates and mineral masses. D,E) Expression of the iron storage genes *ftnA1* and *ftnA2* (D) and the siderophore synthesis genes *asbA* and *asbC* (E). Gene expression was measured by qPCR and normalized to a housekeeping gene *rpoB*. F) Steady state concentration of the siderophore petrobactin, measured by LCMS and normalized within the experiment.

Engineered *A. macleodii* produced siderophores in iron-replete conditions. We continuously cultured the asb+P strain across the same range of reactor conditions we used for the wild type (WT) strain (Fig. S8). Cultures again washed out at high dilution rates, like for the WT strain (Fig. 3C), the washout rate was lower with lower olivine mass (Fig. 3C), and expression of the ferritin genes *ftnA1/2* was lower with low olivine mass and high dilution rates (Fig. 3D). These all suggest that, like for the WT strain, growth was limited by iron in reactors with low olivine mass and high dilution rates. Unlike the WT strain, however, expression of siderophore synthesis genes *asbA* and *asbC* was high across all conditions (Fig 3E). Critically, the highest concentration of petrobactin occurred in the reactor with the highest iron availability (0.2 h^-1^, dilution rate, 0.92 g olivine) indicating that the asb+P strain can produce large quantities of petrobactin even when iron does not limit growth (Fig. 3F).

Engineered *A. macleodii* continuously accelerated olivine dissolution. We operated replicate reactors with the WT and asb+P strains in iron-replete conditions (Fig. 4A). Cell density reached steady state within ∼2 days (Fig. 4B). As expected, the WT strain exhibited high expression of the petrobactin synthesis genes *asbA* and *asbC* only in the absence of olivine, while the asb+P strain exhibited olivine-independent expression of these genes (Fig. 4C). Expression of the ferritin genes *ftnA1/2* was similar between the two strains and indicated that growth was not limited by iron in the presence of olivine. The relative concentration of petrobactin at steady state was >100-fold higher in asb+P cultures than in WT cultures (Fig. 4D). Finally, we measured the total concentration of Ni, Fe, and Si in reactor supernatants at steady state (Fig. S9). Relative to the known stoichiometry of each metal in the olivine substrate, we observed the highest dissolution of Ni, making it the best marker for dissolution at this scale (Fig. 4E). We observed lower dissolution of Si relative to Ni and Fe, especially in reactors with cells. We then compared dissolution rates between reactors (Fig. 4F). The asb+P strain increased the rate of Ni dissolution relative to the abiotic control by a factor of 2.6 ± 1.0 (p = 0.03 after Tukey’s multiple comparison correction). Consistent with our previous observations in these conditions, olivine dissolution with the WT strain was not significantly greater than the abiotic control.

**Fig. 4.**
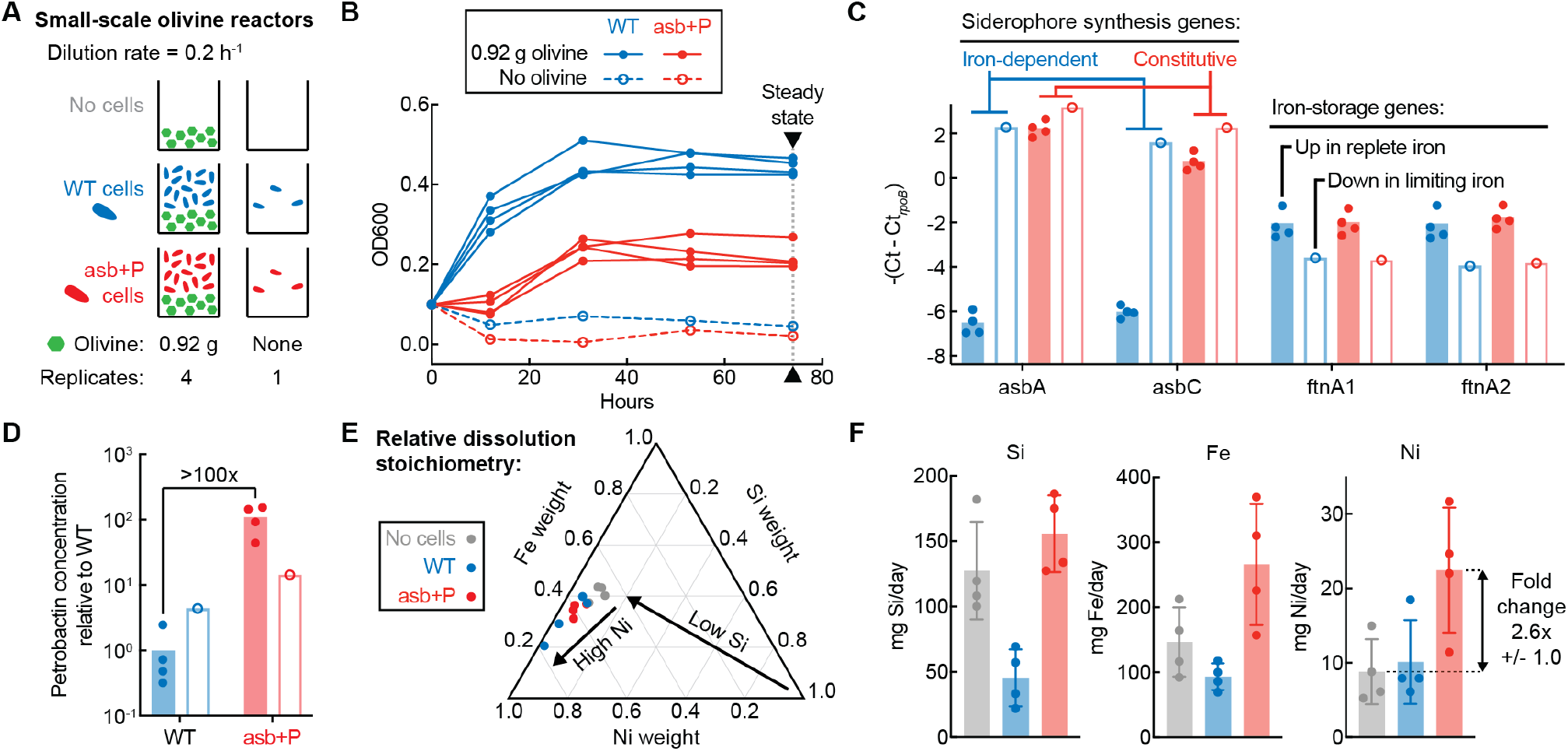
Engineered bacteria continuously accelerate olivine dissolution. A) Experimental layout of parallel mineral bioreactors. B) Optical density of cultures with cells over time after inoculation. Steady state samples were taken after 72 hours of continuous operation. C) Expression of the siderophore synthesis genes *asbA* and *asbC* and the iron storage genes *ftnA1* and *ftnA2* during steady state growth. Gene expression was measured by qPCR and normalized to a housekeeping gene *rpoB*. D) Concentration of the siderophore petrobactin, measured by LCMS and normalized to cultures with the WT strain growing on olivine. E) Relative stoichiometry of mineral dissolution. The concentration of each metal was normalized by the known mass fraction of the metal in the olivine substrate. In the ternary plot, only Ni, Fe, and Si contribute to weighting. F) Dissolution rates of Si, Fe, and Ni from the olivine. Error bars represent 95% confidence intervals calculated by ordinary, 1-way ANOVA.

### Life cycle analysis of large-scale mineral reactors

Since siderophore production can enable continuous acceleration of olivine weathering, we performed LCA to determine the steady state net CO_2_ removal of hypothetical, industrial-scale mineral reactors. We modeled reactors that contain a stationary mineral substrate (>150 tons of olivine sand) submerged beneath a stirred reservoir of seawater that is continuously replaced (Fig. 5A, Table S2). In a control scenario with only seawater flowing over the olivine, we predict gross CO_2_ removal of 0.71 kg/day (Fig. 5B). Emissions associated with pumping energy and olivine procurement are predicted to be only 0.06 kg/day, yielding net CO_2_ removal of 0.65 kg/day.

**Fig. 5.**
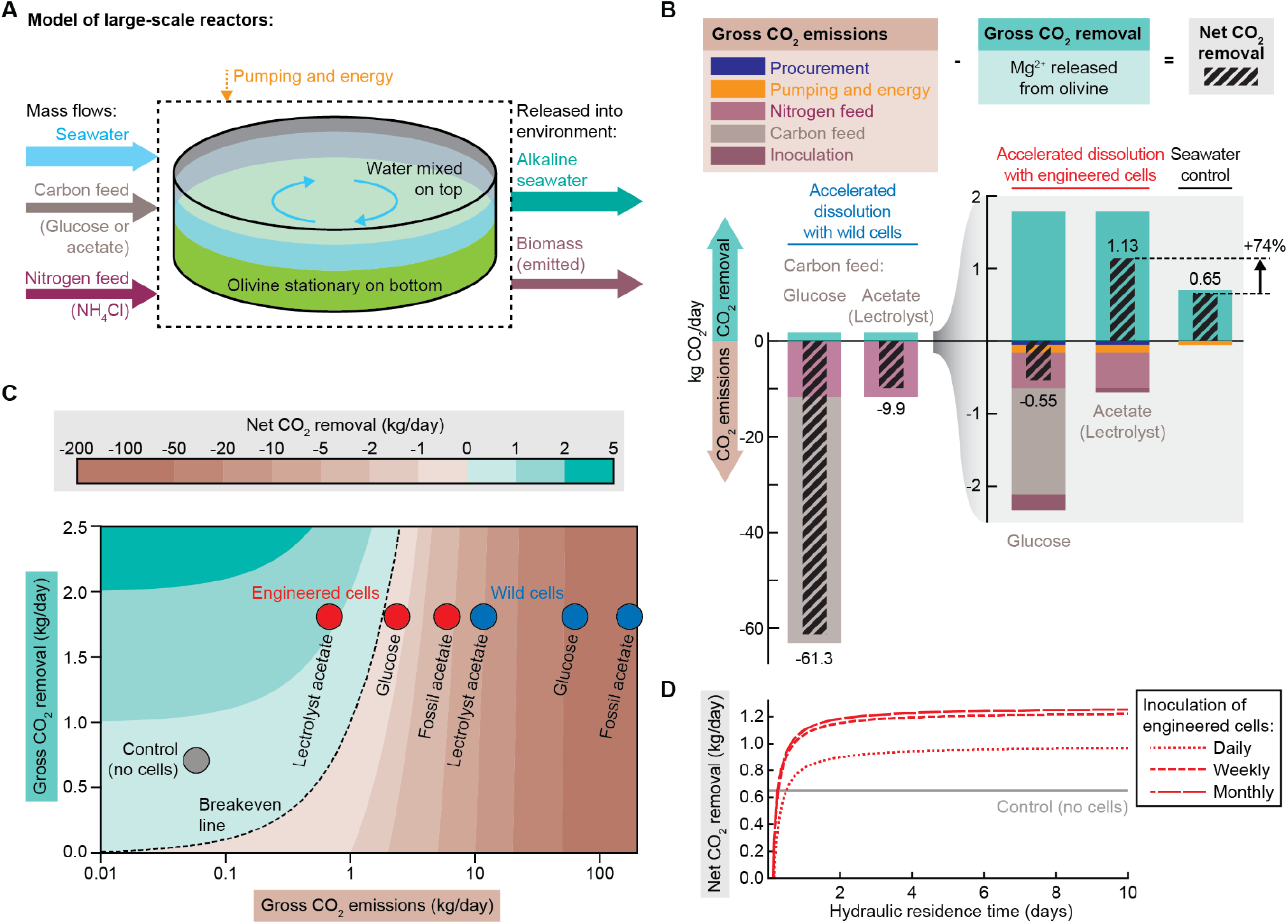
Bio-weathering of olivine can accelerate carbon removal at large scales. A) Schematic of a hypothetical industrial mineral bioreactor. The dotted boundary was used for Life Cycle Analysis of the unit operation. B) Life Cycle Analysis of the bio-weathering unit operation with different carbon feedstocks. The analysis assumes that enough wild or engineered cells are grown to accelerate the olivine dissolution rate. C) Sensitivity analysis of gross CO_2_ emissions compared to gross CO_2_ removal. Choice of carbon feedstock is critical for determining the extent of CO_2_ emissions. D) Sensitivity analysis of the inoculation frequency of engineered bacteria with respect to the hydraulic residence time set by the seawater flow rate.

In a second scenario, nutrients are added to cultivate wild microorganisms naturally present in the seawater to accelerate olivine dissolution. As our small-scale experiments showed, however, wild bacteria only accelerate dissolution during continuous iron-limited growth (Fig. 2, Fig. S5). To achieve continuous iron-limitation, therefore, wild microbes in hypothetical industrial reactors must be fed enough carbon and nitrogen to incorporate all iron released from the olivine into biomass. With this feed rate, we assumed that dissolution could be accelerated by the same fold change that we observed with engineered *A. macleodii* (2.6 ± 1.0, Fig. 4F). With this accelerated dissolution at scale, we predict that the gross CO_2_ removal will increase from 0.71 kg/day to 1.81 kg/day. The required nutrient input, however, would generate CO_2_ emissions of 63.09 kg/day with a glucose feed or 11.71 kg/day with a renewable carbon feed, far exceeding the increase in gross CO_2_ removal (Fig. 5B). Feeding wild microorganisms enough to accelerate olivine dissolution, therefore, would result in significant net CO_2_ emissions.

In a third scenario, engineered *A. macleodii* is periodically inoculated into the reactor and nutrients are added to sustain continuous siderophore production. Since siderophores may be consumed after binding to iron, the quantity of engineered cells required to maximize dissolution of the olivine substrate is not clear. To approximate this nutrient requirement, we calculated the quantity of carbon and nitrogen required to constitute enough petrobactin to solubilize all iron released from the olivine. We assumed that reactors with engineered *A. macleodii* will also increase olivine dissolution by 2.6-fold, resulting in the same gross CO_2_ removal as reactors with wild cells. The nutrient requirement for engineered cells is substantially lower than for wild microorganisms, however, resulting in smaller upstream CO_2_ emissions of only 0.68 kg/day with a renewable feedstock. We predict net CO_2_ removal of 1.13 kg/day in reactors with engineered *A. macleodii*—a 74% increase over the control scenario (Fig. 5B).

According to our LCA, upstream CO_2_ emissions are primarily determined by the carbon feedstock (Fig. 5C). For reactors with engineered *A. macleodii* to outperform control reactors with only seawater, for example, the carbon feedstock must emit less than 0.6 kg CO_2_ per kg C. While we performed our small-scale experiments with glucose as a feedstock, the upstream emissions of a glucose feed (2.8 kg CO2 emitted per kg)^18^ are too high to enable net CO_2_ removal. We also considered acetate, therefore, as a renewable feedstock. While acetate is traditionally produced from fossil fuel combustion, recent technologies enable renewable (CO_2-_ consuming) production of acetate by biological or electrochemical methods. Lectrolyst is a producer of such acetate that uses CO_2_ as a feedstock, removing approximately 1.5 kg CO_2_ per kg acetate.^19^ We conservatively use an emissions factor of 0 kg CO_2_ in our LCA to account for re-emission of biomass from hypothetical large-scale reactors. With this renewable feedstock, reactors with engineered *A. macleodii* fed with acetate achieve net CO_2_ removal and outperform the control scenario (Fig. 5B). In subsequent experiments, we tested the performance of engineered *A. macleodii* when grown on renewable acetate.

Finally, our LCA is sensitive to the frequency of inoculation of engineered *A. macleodii*. Since any pretreatment of the medium is too carbon intensive, engineered cells need to be added directly to unprocessed seawater and survive and function enough to accelerate mineral dissolution. Analysis of the hydraulic residence time and inoculation frequency revealed that engineered cells should function for at least one week (∼7 residence times) to maximize net CO_2_ removal (Fig. 5D). In subsequent experiments, we evaluated the stability and performance of engineered *A. macleodii* in reactors with real seawater.

### Scale-up of biological olivine weathering

To enable scale-up of our bio-weathering process, we created a new engineered strain of *A. macleodii* with enhanced siderophore production that does not require continuous antibiotic selection (Fig. S10A). Instead of placing the *asb* operon on an episomal plasmid, we replaced the native promoter of the *asb* operon in the *A. macleodii* genome with synthetic constitutive promoters. Unlike the plasmid-borne strain, we were able to obtain a strain with the *asb* operon under control of an integrated version of the strongest promoter from our set (J23101). We denoted this strain “asb+G”. In iron replete batch cultures, the asb+G strain produced a similar quantity of petrobactin to the asb+P strain (Fig. S10B).

Next, we sought to test if engineered *A. macleodii* can accelerate CO_2_ removal when grown on a renewable carbon source. To quantify CO_2_ removal, we developed flask-scale mineral reactors that contain 10 g of olivine in ∼150 mL of seawater medium that is continuously replaced (Fig. 6A). Importantly, these reactors use unbuffered medium which allows measurement of the alkalinity of the culture fluid. In theory, dissolution of conservative cations (Mg^2+^) from olivine should confer an increase in the alkalinity of the medium as HCO_3_^-^, a direct measurement of CO_2_ removal from the air (Fig. 1). When we tested flask-scale reactors with a glucose feedstock, our engineered strains maintained high concentrations of the siderophore petrobactin for two weeks (Fig. S11). At pseudo-steady state, however, we observed a decrease in alkalinity (>3 mg CaCO_3_ lost per g glucose fed). We hypothesize that this unexpected reduction was due to the acidic byproducts of glucose metabolism overwhelming the alkalinity from the increase in dissolved olivine.^20^ This biological effect is a second reason why glucose is not a suitable feedstock at industrial scale, in addition to its upstream emissions, as discussed earlier.

**Fig. 6.**
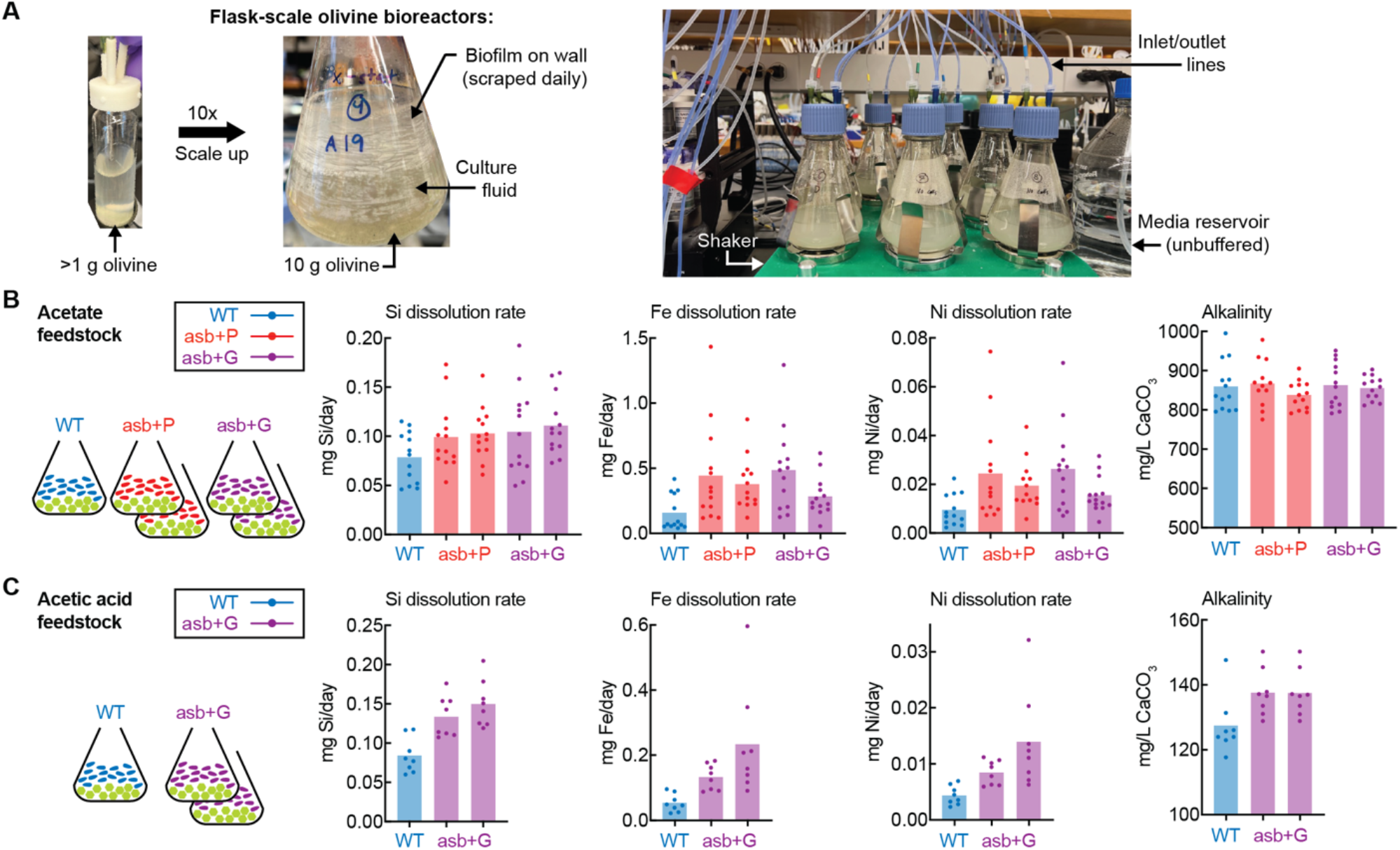
Engineered bacteria continuously accelerate olivine dissolution at flask-scale. A) Flask-scale mineral bioreactors. Unbuffered synthetic seawater media was continuously added and removed at a constant rate while olivine sand remained in the flasks by sinking. Shaking ensures well mixed culture fluid. Biofilms that accumulated on the wall were scraped back into the culture fluid every day. B) Experimental layout of flask reactors fed with acetate (left). Every day, 100 mL of the 150 mL culture volume was removed and replaced with fresh media. Total release rate of three metals from the olivine substrate during pseudo-steady state (middle). Alkalinity of filtered culture supernatants was measured by automatic titration (right). Each bar represents one flask reactor. Replicates represent multiple measurements over time during steady state. C) Experimental layout of flask reactors fed with acetic acid (left). To avoid pH swings, cultures were initiated with acetate. Then, acetic acid media was continuously replaced at 50 mL/day. Measurements at pseudo-steady state were performed as in (B).

To replace glucose, we tested if net alkalinity can be generated in cultures grown on acetate. We confirmed that *A. macleodii* can grow on electrochemically generated acetate solution from Lectrolyst (Fig. S12A). Furthermore, we confirmed that *A. macleodii* did not acidify the culture medium upon consumption of acetate (Fig. S12B). Instead, acetate consumption conferred an increase in the culture pH. In flask-scale experiments, *A. macleodii* was able to continuously grow on acetate (Fig. S13), and engineered strains increased the olivine dissolution rate relative to the WT strain (Fig. 6B).

Despite using a non-acidifying feedstock, however, we still did not observe an increase in the alkalinity of the culture medium. We hypothesize that the high alkalinity of the acetate feed itself masked any differences in alkalinity generated from differences in olivine dissolution. To mitigate this effect and thereby allow us to confirm that differences in olivine dissolution could confer differences in alkalinity, we performed additional flask-scale experiments fed with acetic acid instead of acetate (Fig. S14). Under these conditions, the asb+G strain increased both the olivine dissolution rate and the alkalinity of the culture at steady state (Fig. 6C). While acetic acid is not a viable carbon feed in practice because it lowers the total alkalinity, these experiments confirm that an acetate feedstock, especially if delivered in small quantities, should enable enhanced generation of alkalinity from enhanced mineral dissolution.

Finally, we assessed the stability of engineered *A. macleodii* in unprocessed seawater, since our LCA revealed that engineered strains must survive and function for one week (∼7 residence times) in industrial reactors to maximize net CO_2_ removal. We operated small-scale reactors fed with unprocessed seawater and a concentrated solution of nutrients with acetate as the carbon feed (Fig. S15A). We inoculated separate reactors with the WT strain, the asb+G strain, or no *A. macleodii*. Total growth irrespective of organism was similar in all reactors, demonstrating that organisms present in the seawater feed are also able to consume the nutrient solution (Fig. S15B). We measured the abundance of *A. macleodii* over time by qPCR (Fig. S15C). The WT strain remained abundant for the duration of the experiment (∼20 residence times). While the abundance of the asb+G strain eventually declined, it remained stable for ∼7 residence times. We expect, therefore, that the asb+G strain will survive long enough to only require replenishment once per week in large-scale reactors, thus enabling net CO_2_ removal according to our LCA (Fig. 5D). To test this directly, we constructed pilot-scale mineral bioreactors whose design was guided by these results and those from the flask-scale experiments.

### Pilot-scale mineral bioreactors with real seawater

We demonstrated biologically enhanced olivine dissolution and CO_2_ removal in pilot-scale reactors with real seawater. We placed >4 kg of construction-grade olivine sand in the bottom of 11 L glass reaction vessels (Fig. 7A, S16A). We submerged the sand in ∼2.5 L of unprocessed seawater from the Boston harbor. To best mimic large-scale reactors, we allowed the olivine to remain stationary in each vessel and applied gentle stirring to the ∼1.5 L of water above the sand. To achieve a residence time of ∼1 day in the liquid headspace, we flowed 1.5 L/day of seawater through each vessel.

**Fig. 7.**
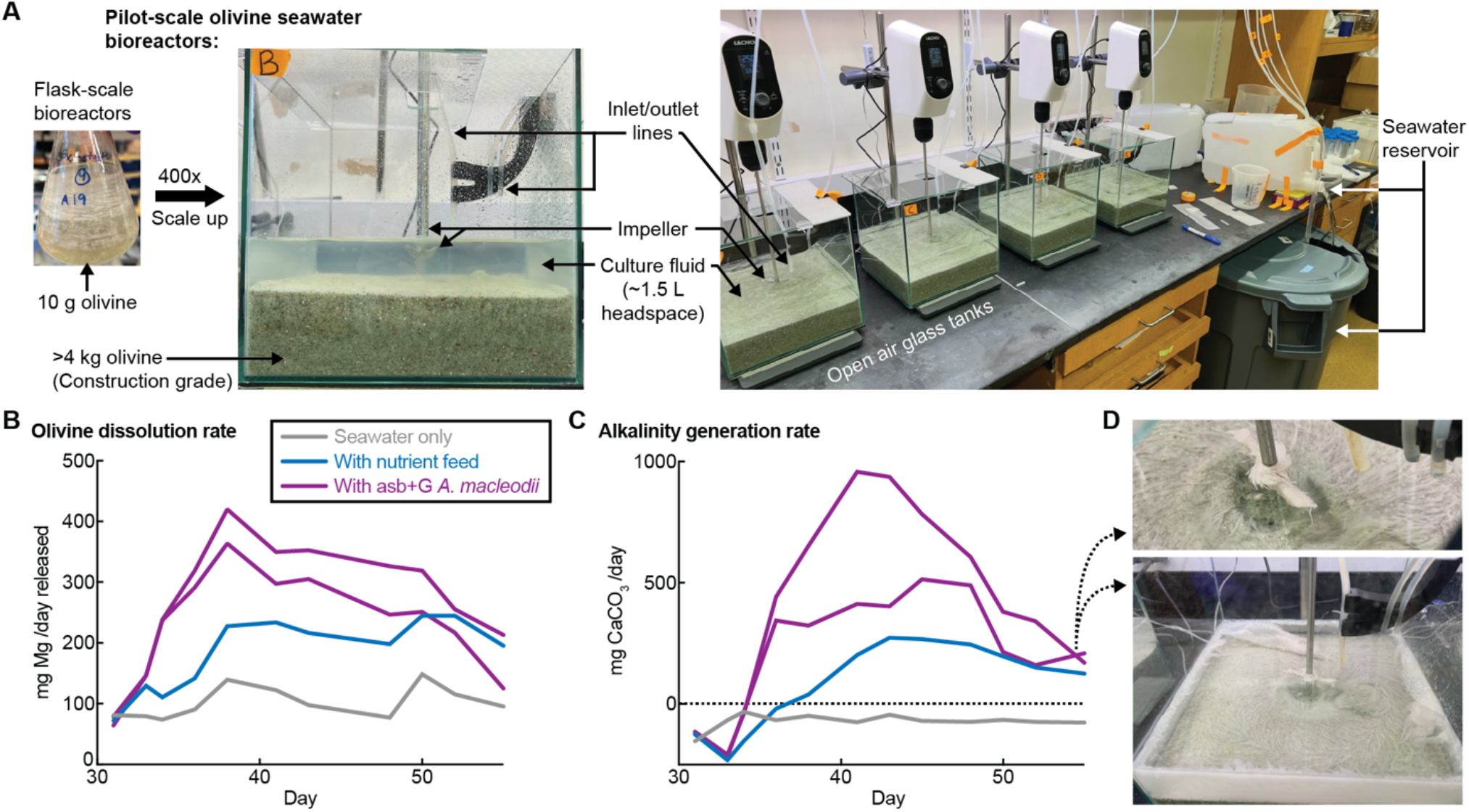
Engineered bacteria continuously accelerate olivine dissolution at pilot-scale. A) Pilot-scale mineral bioreactors. Untreated seawater was continuously added and removed at a rate of 1.5 L/day. Water in the headspace was gently stirred at 200 rpm. Nutrients were continuously added at a 1:50 dilution. Once per week, 140 mL of a saturated culture of the *A. macleodii* asb+G strain was added to the appropriate reactors. B) Total release rate of Mg from the olivine substrate. C) Rate of alkalinity generation in each tank over time, defined as the difference in alkalinity between the culture fluid and the inlet seawater and nutrients, multiplied by the flow rate. D) Representative pictures of biofilms that accumulated on all surfaces in the reactors after ∼3 weeks of weathering.

To determine the rate of nutrient feeding, we operated reactors for one month with only seawater. During this break-in period, the rate of Mg dissolution declined before reaching a pseudo-steady state (Fig. S16B). Using this dissolution rate, we calculated the quantity of siderophores needed to solubilize all the released iron. We then began to continuously add the minimum quantity of nutrients (acetate and ammonium) needed to constitute those siderophores. We also began weekly inoculation of the asb+G strain in two reactors.

In reactors that received the asb+G strain, we observed a 3.1-fold increase (average of days 36-48) in the rate of Mg release from the olivine (Fig. 7B). We also measured the alkalinity of the reactor fluid to assess the amount of dissolved inorganic carbon (HCO_3_^-^) (Fig. S16C). After subtraction of background alkalinity from the inlet media and from the nutrient solution, we observed that reactors with engineered *A*.*macleodii* generated alkalinity at a rate of 0.57 g CaCO_3_ per day (Fig. 7C; average of days 36-48); this is equivalent to 0.50 g CO_2_ per day removed from the air, assuming alkalinity is expressed as HCO_3_^-^.

We observed that seawater ions may interact with the mineral substrate upon dissolution. Ca, specifically, was removed from the seawater in mineral bioreactors both during the break-in period and during the test period (Fig. S16D). Continued removal of Ca from the seawater may explain why the alkalinity in the control reactor was lower than the alkalinity of the inlet seawater at steady state (Fig. 7C). Ca was removed at the highest rate in reactors with engineered *A. macleodii*, suggesting that Ca removal was linked to dissolution of the mineral substrate. The molar ratio of removed Ca to dissolved Mg appeared consistent across all samples (0.64 ± 0.06; Fig. S16E). Interestingly, we observed essentially no dissolution of Si relative to the dissolution of Mg, despite Si being the other major component of the olivine substrate (Fig. S16F). Given that the pH exceeded 9 in all reactors (Fig. S16G), we hypothesize that precipitation of calcium silicate species traps Ca and Si in the reactors.^21^ Indeed, upon completion of the experiment, we noticed that the top layer of the mineral substrate had developed a thin crust.

Finally, after ∼3 weeks of nutrient addition, the rate of Mg dissolution and alkalinity generation declined in all reactors. During this time, we observed significant accumulation of biofilms on the vessel walls and on the top of the olivine sand, particularly in vessels that received inoculations of the *A. macleodii* asb+G strain (Fig. 7D). The biofilms appeared to inhibit mixing of the reactor headspace, and we hypothesize that biofilms act as a physical obstacle to mineral dissolution. Hence, industrial, large-scale reactors will need to be designed to minimize the accumulation of biofilms.

## Discussion

We report here the development of microorganisms with enhanced siderophore production to accelerate alkaline mineral dissolution at scale. First, we demonstrated that since siderophores are only produced during iron-limited growth, continuous production is not feasible in large-scale mineral reactors. To overcome this limitation, we engineered the marine bacterium *A. macleodii* to constitutively produce siderophores. Our strains only contain modifications of the petrobactin synthesis pathway; future efforts to tune the expression of these enzymes or increase the available flux of precursors may further enhance siderophore production.^22^ At the same time, however, engineered strains must retain the ability to grow in competition with the wild seawater microbiome. Even without extensive optimization, our strains conferred a 2.6-fold increase in the rate of olivine dissolution and were sufficiently able to function in unprocessed seawater. Using this data to inform life cycle analysis (LCA), we found that engineered *A. macleodii* can increase net CO_2_ removal by 74% in hypothetical large-scale reactors mineral seawater bioreactors. Any additional increase in the relative rate of dissolution would further increase the predicted rate of gross CO_2_ removal (Fig. 5C).

Our LCA revealed that renewable feedstocks are essential for net CO_2_ removal at scale. We believe that *A. macleodii* is well suited for an industrial seawater process and demonstrated that such a process can achieve net CO_2_ removal when fed a small quantity of an electrochemically synthesized acetate feedstock (Lectrolyst). Siderophores are found in nearly all microbial habitats, however, and a bacterial^23^ or algal^24–27^ host could be engineered for any suitable feedstock and medium.^28^ Indeed, other renewable carbon sources like methanol, waste biomass, or atmospheric CO_2_ may also be suitable nutrients at large scales. Photosynthetic microorganisms that consume CO_2_ also tend to alkalinize their environment, which may confer additional CO_2_ sequestration.^29^

Our LCA also illustrates the need for engineered siderophore production in any microorganism. For wild organisms to continuously produce siderophores, the demand for iron by new biomass must always exceed the iron in the system. Since iron is a micronutrient, this biomass requirement is too high to achieve net CO_2_ removal at scale, even without a carbon feedstock (Fig. 5B). Genetic regulation of siderophore production is broadly conserved across bacteria.^30,31^ In a previous study with batch cultures of the bacterium *Shewanella oneidensis*, for example, olivine dissolution was only accelerated during the short exponential growth phase, likely because siderophore production stopped once cells acquired enough iron to reach stationary phase.^32^ The authors were able to extend the period of enhanced olivine dissolution only by supplementation of the exogenous siderophore desferrioxamine, essentially a manual version of the genetic engineering performed here. This finding may have implications for how microorganisms naturally impact mineral dissolution rates in soil or marine environments. In field trials of Enhanced Rock Weathering, for example, the extent of siderophore production by endogenous bacteria may be limited.^33^

After conceptualization, we constructed and operated pilot-scale mineral bioreactors with continuous flow of real seawater. In reactors that received engineered *A. macleodii*, we observed a 3.1-fold increase in the rate of Mg dissolution from the olivine. This fold change agrees with the fold change measured by dissolution of Ni and Fe at small scales (2.6 ± 1.0). In pilot-scale reactors, this increase in dissolution co-occurred with an increase in the total alkalinity of the culture fluid—a direct measurement of enhanced CO_2_ removal from the air. This also represents permanent CO_2_ storage given the long residence of conservative cations like Mg^2+^ in seawater.^34^

Our control pilot-scale reactor with no nutrient feed did not achieve net CO_2_ removal, possibly due to competing reactions that removed Ca^2+^ ions from the seawater. Secondary precipitation has been widely reported in olivine weathering studies and is expected in tanks that maintain high ion concentrations and high pH.^21,35^ Biologically mediated dissolution, therefore, may represent a strategy to overcome loss of alkalinity in tank-based systems.^6,28^ The reactor with nutrients and only wild organisms was able to generate net alkalinity. We hypothesize that the wild microbiome contained photosynthesizing organisms that alkalinized the culture fluid upon consumption of NH_4_Cl and CO_2_. The additional generation of alkalinity upon addition of engineered *A. macleodii*, therefore, was due to accelerated olivine dissolution from removal of iron by siderophores.

In our pilot-scale systems, most dissolution likely occurs only from the top layer (<1 cm deep) of the stationary mineral substrate. Both the thin layer of hardened crust at the end of the pilot-scale experiment and calculations of the surface area normalized dissolution rate support this assumption (Table S2). At scale, periodic mechanical disruption of the top layer of the mineral substrate may alleviate passivation from both secondary mineral precipitation or from biofilm formation and enable accelerated weathering on longer time scales (>1 month). Biofilm formation is common in many industrial processes with unprocessed seawater, and biofouling removal is common practice in wastewater and bioleaching operations.^36,37^ Furthermore, while we modeled large-scale reactors that contain ∼180 tons of olivine in a one-meter-deep pile, we assumed that the mineral dissolution rate would scale with the area of the tank basin—that is, the interfacial area between the mineral pile and the stirred seawater. In practice, therefore, large-scale reactors could contain a much smaller quantity of olivine that is replenished more frequently, which may prevent mass transport limitations within the mineral heap.

In summary, we report here a new unit operation for microbial mineral processing. This unit operation is distinct from recent trends in fermentation technology towards intensified bioprocesses—our strategy is to feed the minimal quantity of nutrients required to accelerate mineral dissolution. With no control over temperature or sterility, mineral bioreactors more closely resemble operations for water treatment or mineral heap bioleaching. The deployment of engineered biocatalysts in an unprocessed, ‘dirty’ seawater medium will allow direct deployment into the existing Enhanced Rock Weathering industry and will enable development of new environmental bioprocesses.

## Acknowledgements

LCMS was performed at the Harvard Center for Mass Spectrometry – A Harvard FAS Division of Science Core Facility; we thank Jennifer Wang. ICP-OES was performed at the MIT Nano Core; we thank Tim McClure and Shayne Harrelson. Plasmids for transformation of *A. macleodii* were kindly provided by Chris Dupont and Erin Garza. Seawater for pilot-scale systems was acquired at the New England Aquarium; we thank Jackie Anderson. We thank Martin Van Den Berghe for scientific guidance and geochemical methods. We thank Nate Walworth for scientific guidance and business context for mineral dissolution. We thank Ethan Jones and Simon d’Oelsnitz for guidance on operon design.

## Author contributions

N.C.D., A.P.J., P.A.S., and M.S. conceived the study. N.C.D. and A.P.J. constructed the theoretical model. N.C.D., M.H., and A.P.J. constructed and operated chemostats and bioreactors. N.C.D. and J.T.B. constructed and characterized bacterial strains. N.C.D. performed metals analysis. J.T.B. performed analysis of gene expression. A. F. and N.C.D. modeled industrial reactors and performed Life Cycle Analysis. Q.A.J., S.J.D., P.A.S., and M.S. advised on experimental design and analysis. N.C.D. wrote the manuscript. All authors edited the manuscript.

## Data and materials availability

All data are available in the manuscript or the supplementary materials. Code used to generate the chemostat model can be found at https://github.com/amoghpj/rock-weathering.

Custom code for operation of eVOLVER chemostats can be found at https://github.com/amoghpj/evolver/releases/tag/V0.1.

