## Supplementary material for "Engineered bacterial siderophore production accelerates rock weathering for carbon removal": 20260506 supplemental materials.docx

**Continuous accelerated rock weathering by marine bacteria with enhanced siderophore production**

Supplemental materials

Methods and materials

Table S1. Composition of olivine sand

Fig. S1. Refence gene expression in *A. macleodii*

Fig. S2. Batch growth of *A. macleodii*

Fig. S3. Continuous growth of *A. macleodii* on olivine sand

Fig. S4. Measurement of olivine dissolution in cell culture

Fig. S5. Mineral chemostats with wild type bacteria

Fig. S6. Characterization of synthetic promoters in *A. macleodii*

Fig. S7. CAS agar assay of strains carrying plasmids

Fig. S8. Mineral chemostats with the asb+P strain

Fig. S9. Comparison of olivine dissolution rates

Fig. S10. Engineering of *A. macleodii* for stable siderophore expression

Fig. S11. Flask-scale weathering with glucose feed

Fig. S12. Batch growth of *A. macleodii* on acetate

Fig. S13. Flask-scale weathering with acetate feed

Fig. S14. Flask-scale weathering with acetic acid feed

Fig. S15. Growth of *A. macleodii* in unprocessed seawater

Fig. S16. Pilot-scale bio-weathering

Appendix 1: Model of mineral chemostats

References

Methods and materials

*Bacterial strains and media*

Wild type *Alteromonas macleodii* was acquired from the ATCC (#27126). The asb operon was amplified from the *A. macleodii* genome by PCR (NEB Q5® High-Fidelity 2X Master Mix) and assembled using Gibson assembly (NEBuilder® HiFi DNA Assembly). Plasmids were cloned and propagated in *Escherichia coli* DH5-alpha (NEB). *A. macleodii* was transformed by conjugation from an *E. coli* vector strain as described previously.^1^ Multiple transformants were sequence verified and stored as frozen glycerol stocks. Sequences to enable recombination of the KanR sequence for disruption or upregulation of the asb locus were also amplified from the genome and cloned by Gibson assembly. For genomic modification of the ∆asbAB and asb+G strains, plasmids were transformed by conjugation and transformants were grown on sucrose medium as described previously.^2^ A stable clone with genomic modification was isolated and verified by PCR amplification and Sanger sequencing. Sequences of all plasmids used are included. Cells were retrieved from frozen stocks by plating for single colonies on complex media (BD Difco Marine Broth 2216) with 1.5% agar at 37°C. Strains carrying plasmids were cultivated with 25 µg/L kanamycin as needed, added from a 1000x stock. To remove excess iron, kanamycin was stirred for 24 h with 50 g/L Chelex100 resin (Bio-Rad) and filtered before use.

Synthetic medium was prepared by combining 36 g/L synthetic seawater (Instant Ocean), 0.756 g/L NH_4_Cl, 0.127 g/L K_2_HPO_4_. Experiments at small-scale used media buffered with MOPS (11 g/L for batch experiments, 2.75 g/L for continuous experiments), followed by adjustment to pH 7.4 with NaOH, and filtration. MOPS buffer has been previously shown to have limited interaction with cations released from olivine.^3^ Glucose was prepared as a 200 g/L solution, filtered, and added immediately prior to use to a final concentration of 3.6 g/L for batch experiments. Continuous and semi-continuous experiments were performed with 0.9 g/L glucose such that cell yield with excess iron is limited by glucose (Fig. S1B). Trace metals (ZnSO_4_-7H_2_O 220 µg/L, MnCl_2_-4H_2_O 1.8 mg/L, CoCl_2_-6H_2_O 400 µg/L, CuSO_4_-5H_2_O 80 µg/L, NaMoO_4_-2H_2_O 2.4 mg/L, Na_2_SeO_3_ 2 µg/L, H_3_BO_3_ 3 mg/L, NiCl_2_-6H_2_O 4 µg/L) and vitamins (cyanocobalamin 0.05 µg/L, biotin 0.05 µg/L, thiamine HCl 100 µg/L) were also added immediately prior to use from 1000x stocks. Cultures that required additional soluble iron were supplemented with 0.81 mg/L FeCl_3_ from a 1000x stock.

Olivine composition is listed in Table S1. Pure olivine sand (AFS50, Sibelco) was used in small-scale and flask-scale experiments. Before small-scale experiments, sand was pretreated by several days of shaking in deionized water, one day of shaking in ethanol, and drying at 80°C. Sand was dispensed into culture vials using a customized solids dispenser (XQ Instruments). For flask-scale experiments, sand was not pretreated. Construction grade olivine sand (LE30, AGSCO) was used in pilot reactors. Before loading into reactors, silt was removed from the olivine by serial replacement of deionized water in a concrete mixer.

All media was prepared in plasticware that was rinsed 3x with deionized water to remove excess metals. Glass media bottles and culture vials for continuous experiments were washed 3x with 1% HNO_3_ and 3x with deionized water before autoclaving.

*Chrome azurol-S (CAS) assay*

CAS dye was prepared as a 10x concentrate as described previously^4^ and mixed into synthetic minimal medium with 1.5% agar. Transformant colonies were grown in 5 mL culture tubes in marine broth for 24 hours at at 250 rpm at 37° C, and 30 µL of each culture was plated onto the center of a CAS agar plate. Plates were stored in the dark at room temperature and photographed intermittently for two weeks.

*Batch cultures*

Cells were streaked from frozen glycerol stocks onto solid agar marine broth medium for 1 day at 37°C. A single colony was picked and grown in a 5 mL culture in marine broth in a plastic test tube for 1 day. Saturated cultures were added at a 1:1000 dilution to either 50 mL of synthetic medium in a 250mL plastic flask or 4 mL of synthetic medium in a 24-well plastic plate. Tubes, plates, and flasks were shaken at at 225 rpm at 37°C. Optical density at 600 nm (OD600) was measured by sampling 1 mL of culture into a cuvette and reading on a spectrophotometer. Culture supernatant was harvested by centrifugation for 15 minutes at 12,000xg followed by filtration and storage at 4°C.

*Continuous cultures*

Small-scale chemostats were based on eVOLVER chemostats,^5^ operated with fresh type 1 borosilicate glass vials (Chemglass), plastic stir bars (Fisherbrand Octagon Spinbar Magnetic Stirring Bars), and straight, blunt end needles (Fisnar QuantX) fitted into custom vial caps. Vials were loaded with olivine sand before inoculation according to the experimental design. Chemostats were operated at 37° C with stir speed 8. Dilution rate was specified by the experimental design and controlled with a custom software wrapper.

For flask scale experiments, media was continuously added and removed via blunt end needles that were inserted into plastic flasks. Needles were placed to maintain a culture volume of ~150 mL. Flow was controlled with eVOLVER pump arrays, and flasks were shaken at 225 rpm.

At small- and flask-scale, continuous experiments were inoculated with batch flask cultures such that the initial OD600 = 0.1. Media was prepared in advance and drawn from a glass bottle reservoir. Optical density at 600 nm (OD600) was measured by sampling 1 mL of culture into a cuvette and reading on a spectrophotometer.

Pilot-scale mineral reactors were constructed from glass tanks (3-gallon Imagitarium fish tanks, Petco) and overhead stirrers (Lachoi 20L overhead stirrer, Amazon). Seawater was obtained monthly from the Boston harbor and stored at 4° C in the dark before being added to a 40-gallon reservoir with a circulation pump. During operation, seawater was added to the reactors at a rate of 1.5 L/day in 1 mL increments. The nutrient solution (34.26 g/L CH_3_COOK, 6.59 g/L NH_4_Cl, 1.11 g/L K_2_HPO_4_) was added at a rate of 30 mL/day in 1 mL increments. Culture fluid was continuously removed through an outlet line that was placed such that the liquid headspace was ~1.5 L. To initiate reactors, construction grade olivine was washed, dried, added to vessels, and leveled. Vessels were filled by the pumps over several days as to not disturb the olivine. Stirrers were set to 200 rpm as to not disturb the olivine. Samples were drawn by pipette from the culture fluid near the outlet line.

*Quantitative PCR*

To measure gene expression, 5-10 mL of culture fluid was sampled and centrifuged for 15 minutes at 3,000 x g at 4° C. After decanting of supernatant, pellets were flash frozen and stored at -80° C. After storage, pellets were resuspended in 200 µL of Max Bacterial Enhancement reagent (Thermo), moved to an RNAse-free microcentrifuge tube, and incubated in a heat block at 94° C for 5 minutes. The heated sample was then lysed by addition of 1 mL of TRIzol Reagent (Thermo) and mixed by pipetting. RNA was then extracted using the Direct Zol RNA Miniprep Kit (Zymo) according to the manufacturer’s instructions. RNA was eluted from the columns in nuclease free water and stored at -80° C until further processing.

Following RNA extraction, quantitative PCR (qPCR) was performed with the Luna Universal One-Step RT-qPCR Kit (NEB) according to the manufacturer’s instruction. Each unique primer-RNA combination was performed in triplicate 20 µL reactions in a 384-well plate. RT-qPCR was conducted using the QuantStudio 7 Flex (Applied Biosystems by Thermo Fisher Scientific). Gene expression is reported as the difference in Ct value relative to the reporter gene *rpoB*.

In experiments with untreated seawater, the abundance of the *A. macleodii* genome over time was measured by TaqMan Probe based qPCR. Culture samples were stored frozen at -80 until they were processed for qPCR. Samples were lysed by mixing 1:1 into 40 mM NaOH and boiling at 95° C for 10 minutes. The lysed cultures were then diluted 1:200 into sterile water to dilute out the salt from the sea water. 2 uL of the lysed and diluted sample was used as input to the qPCR reaction. *A. macleodii* specific primers and probes were designed using primergen (<https://github.com/amoghpj/primergen>). (MP154:A_macl_F TGAGACAGAACTGCAGCGTT, MP154:A_macl_R GGCTGCACCGTGGTATCG, MP154:A_macl_P CGTGGTATCGCCGATAACTAGCGC). Genome copy numbers per mL were calculated from Cts as follows:

$$1000 \frac{\mu L}{mL}*200 \left( dilution \right)*\frac{2^{35-Ct}}{2 \mu L input}$$

There was no detected *A. macleodii* signal from seawater controls within limit of detection.

*Mass spectrometry of petrobactin*

To measure relative petrobactin concentration, 5 mL of culture fluid was sampled and centrifuged for 10 minutes and 3,000 x g. Supernatant was filtered and stored at 4° C if needed. Samples were prepared for LCMS on 6 mL Bond Elut ENV columns (Agilent). All washes were performed by gravity at room temperature. New columns were washed 3x with 4 mL of methanol, then 3x with 4 mL of 0.01 M hydrochloric acid. Columns were then loaded with 4 mL of culture supernatant, and washed 2x with deionized water. To elute, columns were washed 3x with 4 mL methanol and the effluent was collected into clean tubes. Effluents were then concentrated by evaporation by Speed-Vac (Thermo) to a volume of 1-3 mL. Finally, 400 µL of concentrated effluent was centrifuged for 20 minutes at 20,000 x g and transferred to a clean vial. LCMS was performed on a Thermo Scientific Dionex UltiMate 3000 UHPLC coupled to a Thermo Q Exactive Plus mass spectrometer system equipped with an HESI-II electrospray ionization source. Data were acquired with Chromeleon Xpress software for UHPLC and Thermo Xcalibur software for mass spectrometry (version 3.0.63), and processed with Thermo Compound Discoverer software (version 3.3 SP3). Petrobactin was identified by its mass as described previously.^2^

*Flow cytometry*

To quantify the strength of synthetic promoters J23100, J23101,J23105, J23107, J23112, J23113 and J23115 in *A. macleodii*, the intensity of GFP fluorescence in the different strains of *A. macleodii* was measured via flow cytometry. Frozen stocks of *A. macleodii* strains carrying plasmids for expression of GFP were streaked for single colonies on agar marine broth medium and incubated at 37 °C over night. Three colonies of each strain were picked for biological triplicates. Each colony was used to inoculate two 3 mL liquid precultures in synthetic medium, one with 5 µM FeCl_3_ and one with no additional iron. Precultures were grown at 37 °C at 225 rpm overnight. Precultures were used to inoculate 5 mL cultures with the same media, which were incubated at 37 °C at 225 rpm for 6 h. After incubation, 20 µL of culture was mixed by pipetting, filtered through a 50 μL filter to remove aggregates, and diluted 1:10 in PBS. Cells were then run on a S1000EON Flow Cytometer (Stratedigm). Fluorescence was measured for FITC (530/30) after excitation at 488 nm with a flow rate of 0.5 mL/min. The FSC and SSC were set to 50, the FITC -sensitivity was set to 3%. Each readout was stopped after 20,000 events or a runtime of 120 sec. A wild type control was used to select the gating settings for all samples. Data was analyzed using the FlowCal library in python 3.11. Each replicate reported is the geometric mean of fluorescence.

*Inductively coupled plasma optimal emission spectroscopy (ICP-OES)*

For metals analysis, 5 mL of culture was sampled from continuous culture vials into 15 mL tubes (Falcon). Samples were acidified with 150 µL of 70% HNO_3_, then centrifuged for 10 minutes at 3,000 x g. Supernatant was decanted into temporary plastic storage tubes and stored overnight at room temperature. Cell pellets were further acidified with 200 µL of 70% HNO_3_ and resuspended by pipetting. Resuspended pellets were allowed to digest at 80° C overnight. Supernatants were added back into the original tube after digestion of the pellet. Combined samples were passed through a 0.2 µm filter. Samples were run on an Agilent ICP-OES 5100 with ICP Expert software (version 7.3.0.8799) and compared against a multiple element standard curve (High Purity Standards). Metal lines (Fe 238.204, Ni 231.604, Si 251.611) were analyzed for concentration. To calculate the rate of release of each metal from the olivine substrate, the concentration of the metal in control vials without olivine was subtracted from the vial of interested. The resulting concentration was multiplied by the appropriate dilution rate and normalized if necessary.

*Alkalinity assay*

Culture fluid from extended weathering experiments was harvested, centrifuged for 20 minutes at 12,000 x g, and filtered. Filtered supernatant was loaded into a mini titrator (Hanna Instruments HI84531U-01) and analyzed in the low-range according to the manufacturer’s instructions.

*Life-cycle assessment (LCA) modeling*

A comparative LCA was developed to quantify the operational net CO₂ sequestration potential of the continuous reactor systems. Net CO₂ flux was defined as the gross CO₂ captured via the measured stoichiometric magnesium (Mg) release minus the summed upstream greenhouse gas (GHG) emissions. The assessed system boundary encompassed olivine procurement (mining and comminution), operational electricity for continuous fluid pumping and stirring, periodic microbial inoculation, and the continuous supply of carbon and nitrogen nutrient feeds.

Olivine comminution emissions were dynamically calculated using an empirical power-law model derived from Renforth et al.^6^:

$$Emissions = 165.01 * d^(-0.377)$$

corresponding to the empirical particle size distribution (d) of the experimental Sibelco AFS50 sand. Operational kinetic energy demands for stirring and pumping were converted to GHG totals using a conservative estimate of localized Massachusetts grid emissions factor of 0.482 kg CO₂e/kWh. Upstream emissions burdens for carbon nutrient sources were evaluated across multiple programmatic scenarios: proxy fossil-derived potassium acetate (calculated at 2.11 kg CO₂e/kg product, derived proportionally from standard acetic acid [1.87 kg CO₂e/kg]^7^ and potassium hydroxide [1.25 kg CO₂e/kg]^8^ manufacturing impacts) with a 0.25 kg CO_2_e/kg estimated drying tax, standard industrial glucose (1.10 kg CO₂e/kg)^9^, and a theoretical carbon-neutral electrochemical acetate (0.0 kg CO₂e/kg). Nitrogen supplementation was modeled uniformly as ammonium chloride with an emission factor of 4.50 kg CO₂e/kg N.^10^ Total daily modeled emissions were normalized against the experimental daily Mg mass flux to output the net removal rates across varying biological configurations and hydraulic residence times.

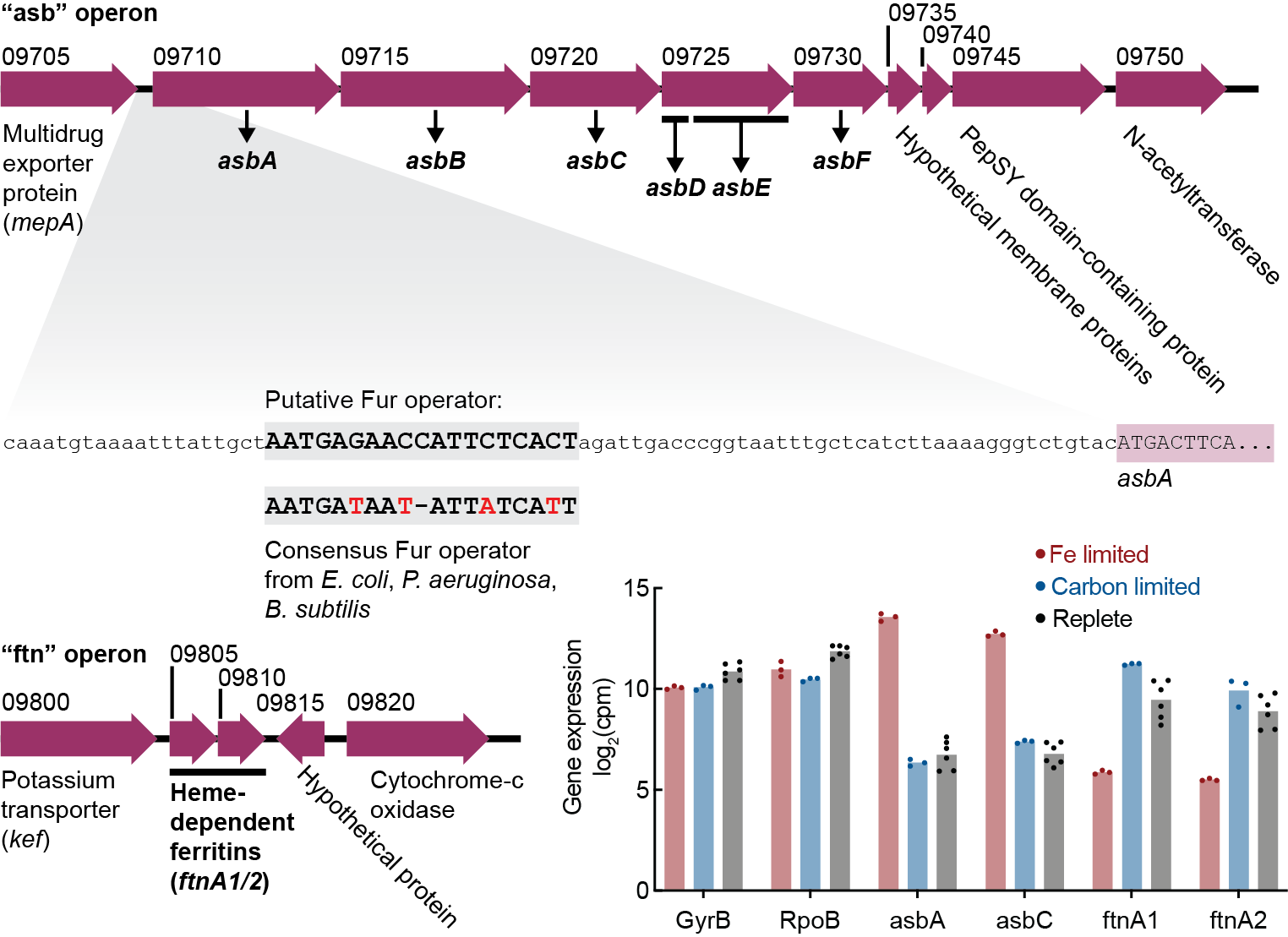

**Fig. S1. Refence gene expression in *A. macleodii*.** Diagram of the *asb* operon for synthesis of the siderophore petrobactin and the *ftn* operon for expression of ferritin proteins. The consensus Fur operator was identified by visual inspection. Gene expression data was adapted from Manck et al.^11^ *gyrB* and *rpoB* were selected as housekeeping genes that are expected to maintain constant expression levels.

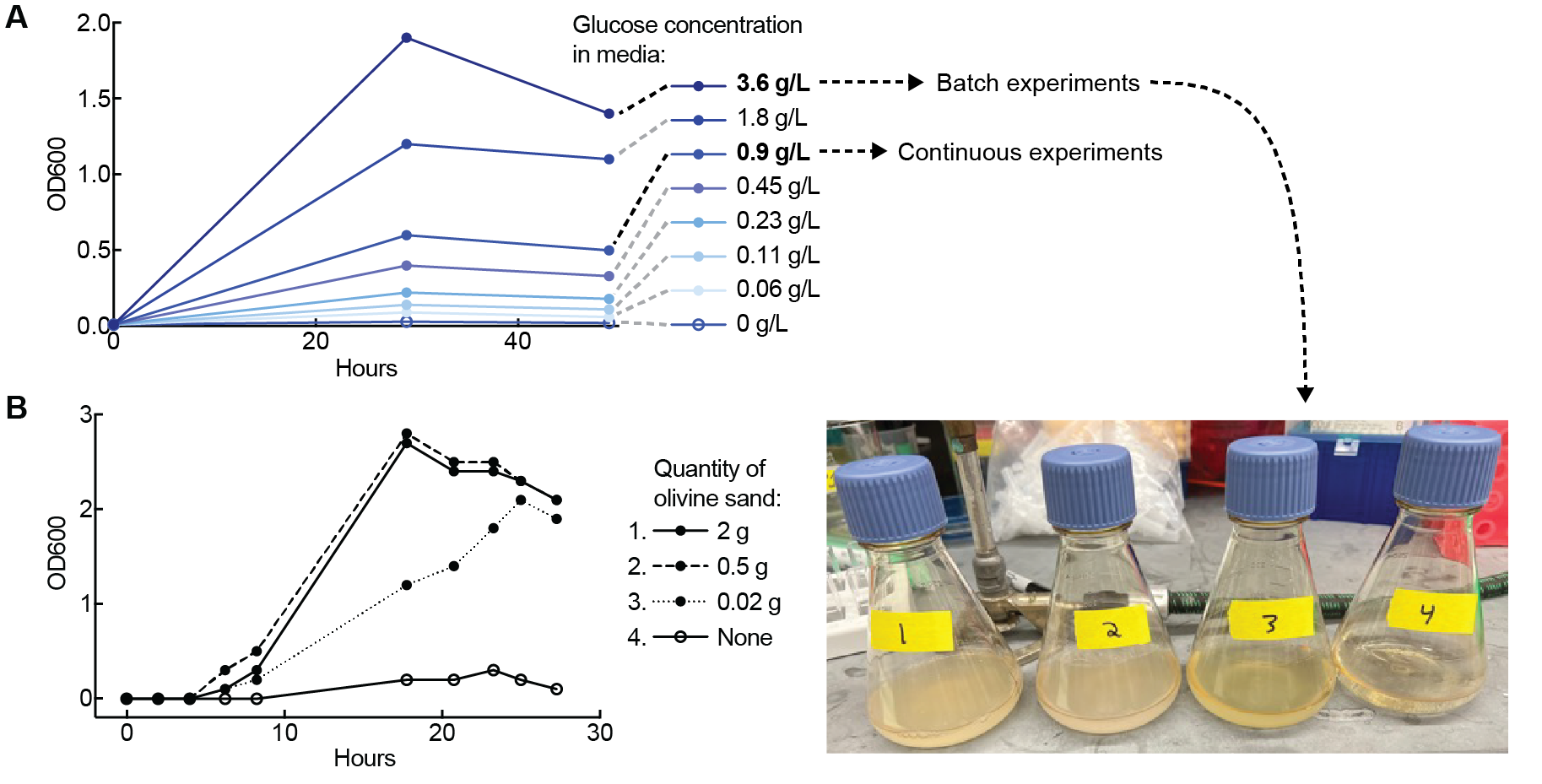

**Fig. S2. Batch growth of *A. macleodii***. A) Growth of 50 mL shake flasks measured by optical density at 600 nm. Flasks contained buffered seawater medium with replete iron (5 µM) and varying quantities of glucose. A glucose concentration was selected for subsequent continuous cultures such that the cell yield in iron-replete conditions would be limited by glucose. B) Growth of 50 mL shake flasks measured by optical density at 600 nm. Flasks contained buffered medium, 3.6 g/L glucose, and no soluble iron. Flasks are depicted in the photograph on the right.

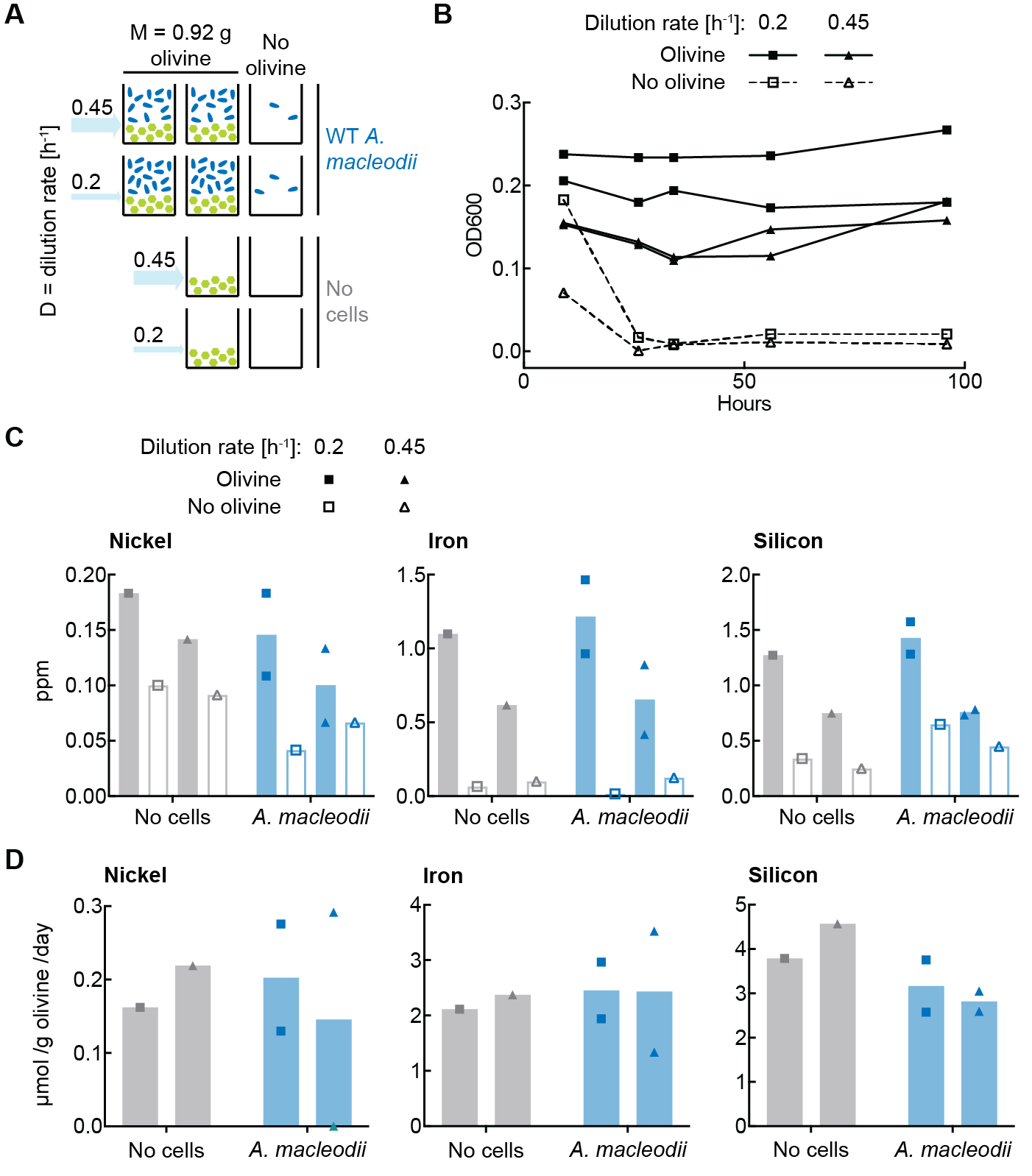

**Fig. S3. Continuous growth of *A. macleodii* on olivine sand.** A) Experimental layout of parallel chemostats. Two dilution rates were tested. B) Optical density of cultures with cells over time. Steady state samples were taken after 56 hours of continuous operation. By 96 hours, biofilms were visible on the vessel walls. C) Steady state total concentration of three metals in the culture fluid, measured by ICP-OES. D) Steady state total release rate of three metals from the olivine substrate.

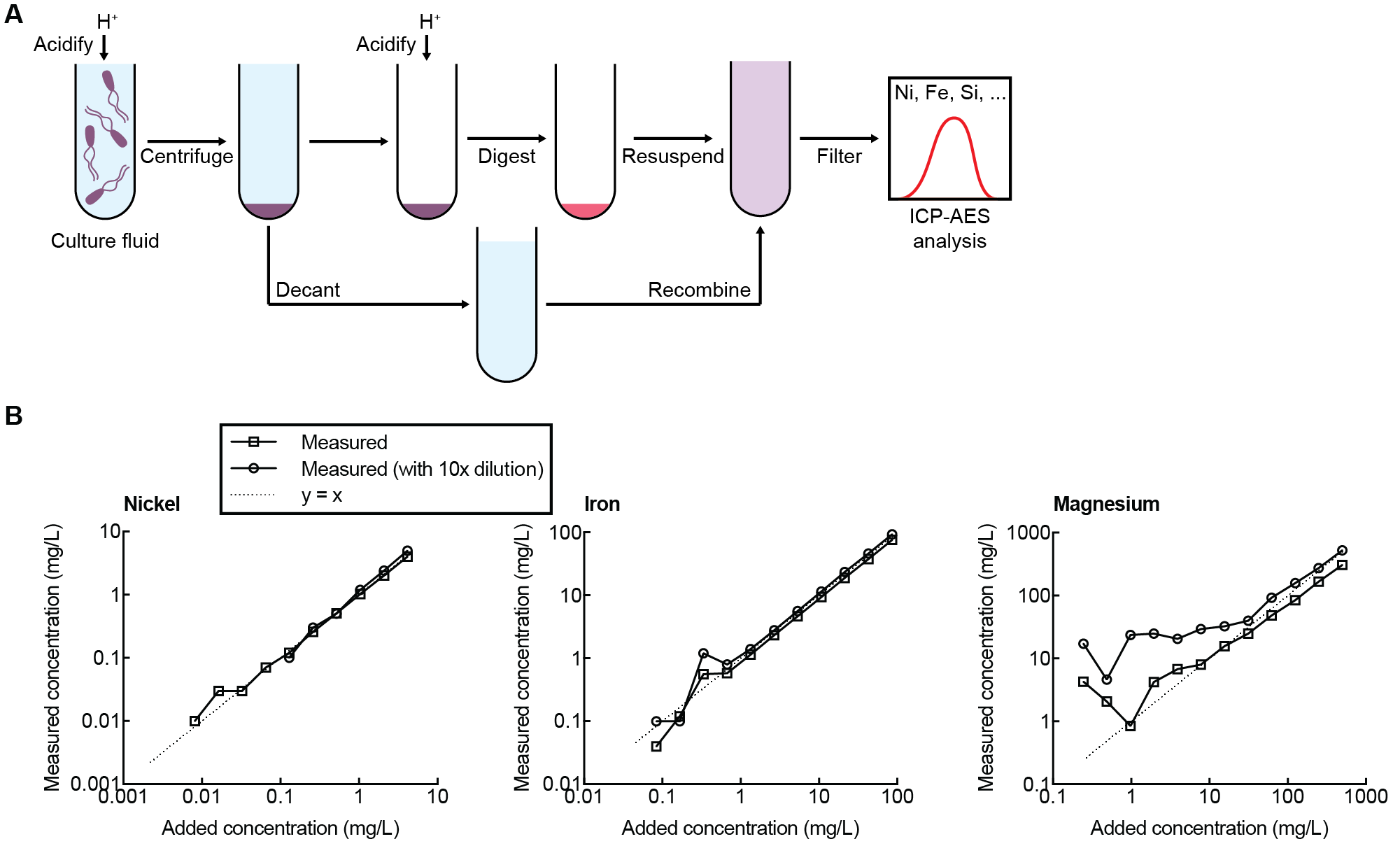

**Fig. S4. Measurement of olivine dissolution in cell culture.** A) Schematic of culture fluid preparation for spectroscopic analysis. Extracellular and cellular fractions were separated for digestion in acid and recombined for measurement by ICP-OES. B) The sample preparation method was tested by adding a known concentration of metals to *A. macleodii* cell culture. Metals were added all together, in ratios that represent the composition of olivine. The sample was then prepared as shown in (A) and measured, with and without a 10x dilution.

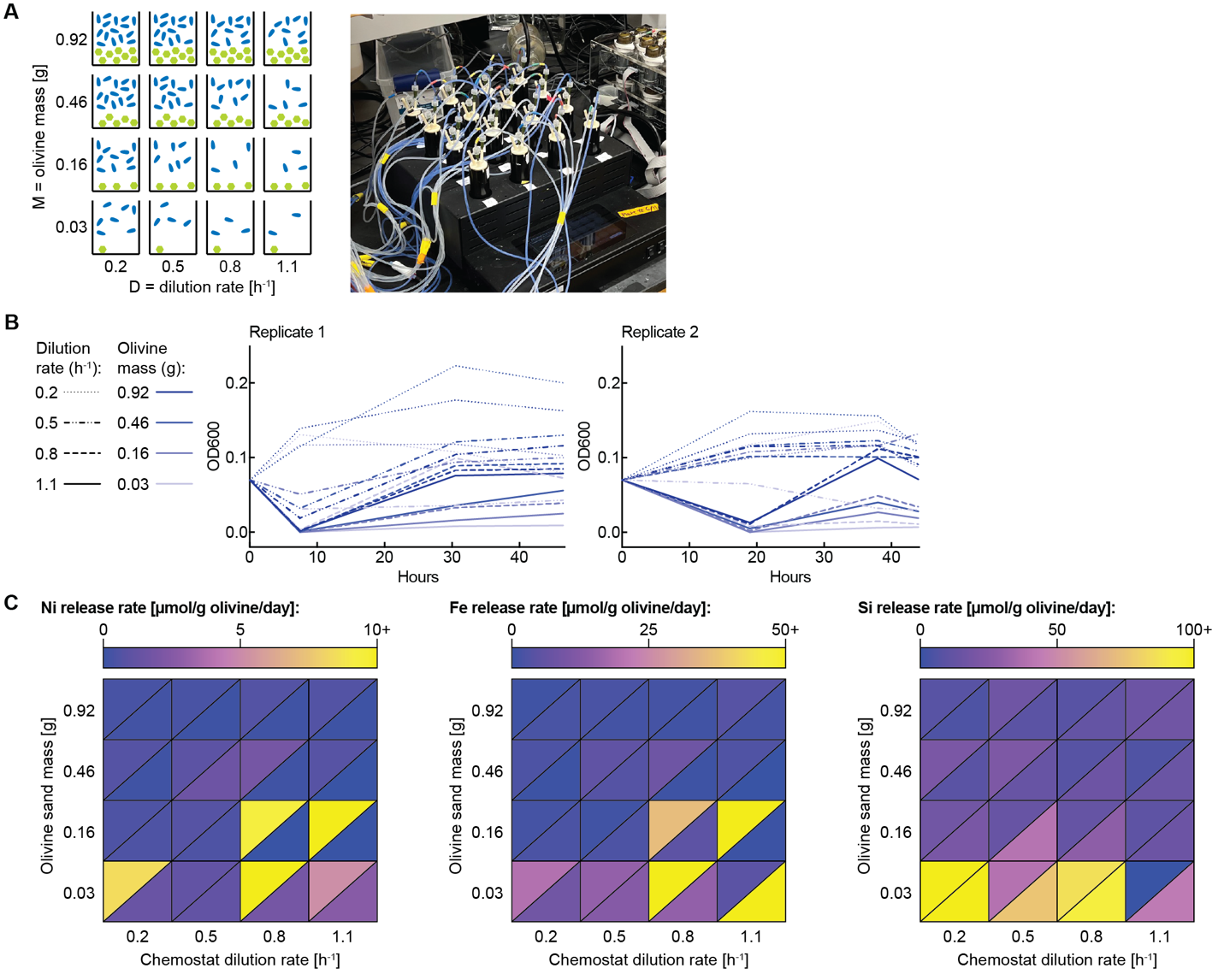

**Fig. S5. Mineral chemostats with wild type bacteria.** A) Experimental layout of parallel chemostats. B) Optical density of cultures over time after inoculation. C) Steady state total release rate of three metals from the olivine substrate, measured by ICP-OES.

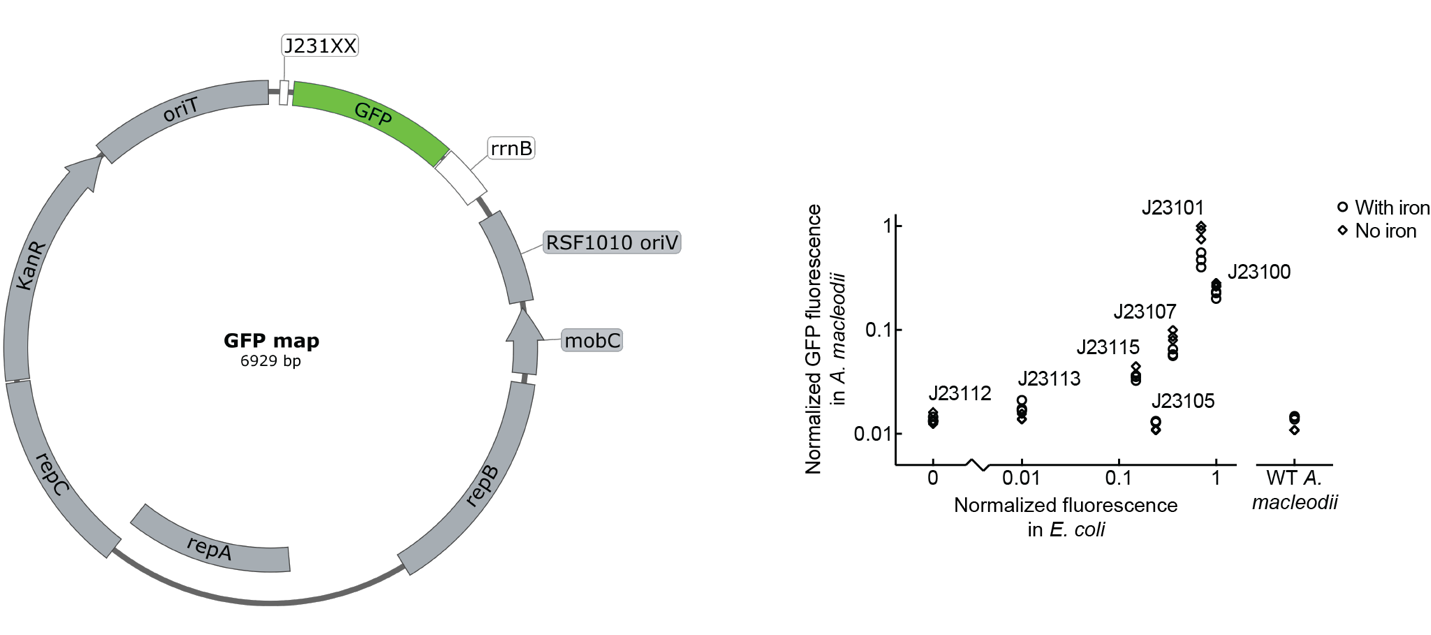

**Fig. S6. Characterization of synthetic promoters in *A. macleodii*.** Cells were transformed with plasmids expressing GFP with one of the synthetic promoters, cultivated in 5 mL of complex media with 5 µM iron or without iron, and GFP expression was measured by fluorescence using flow cytometry.

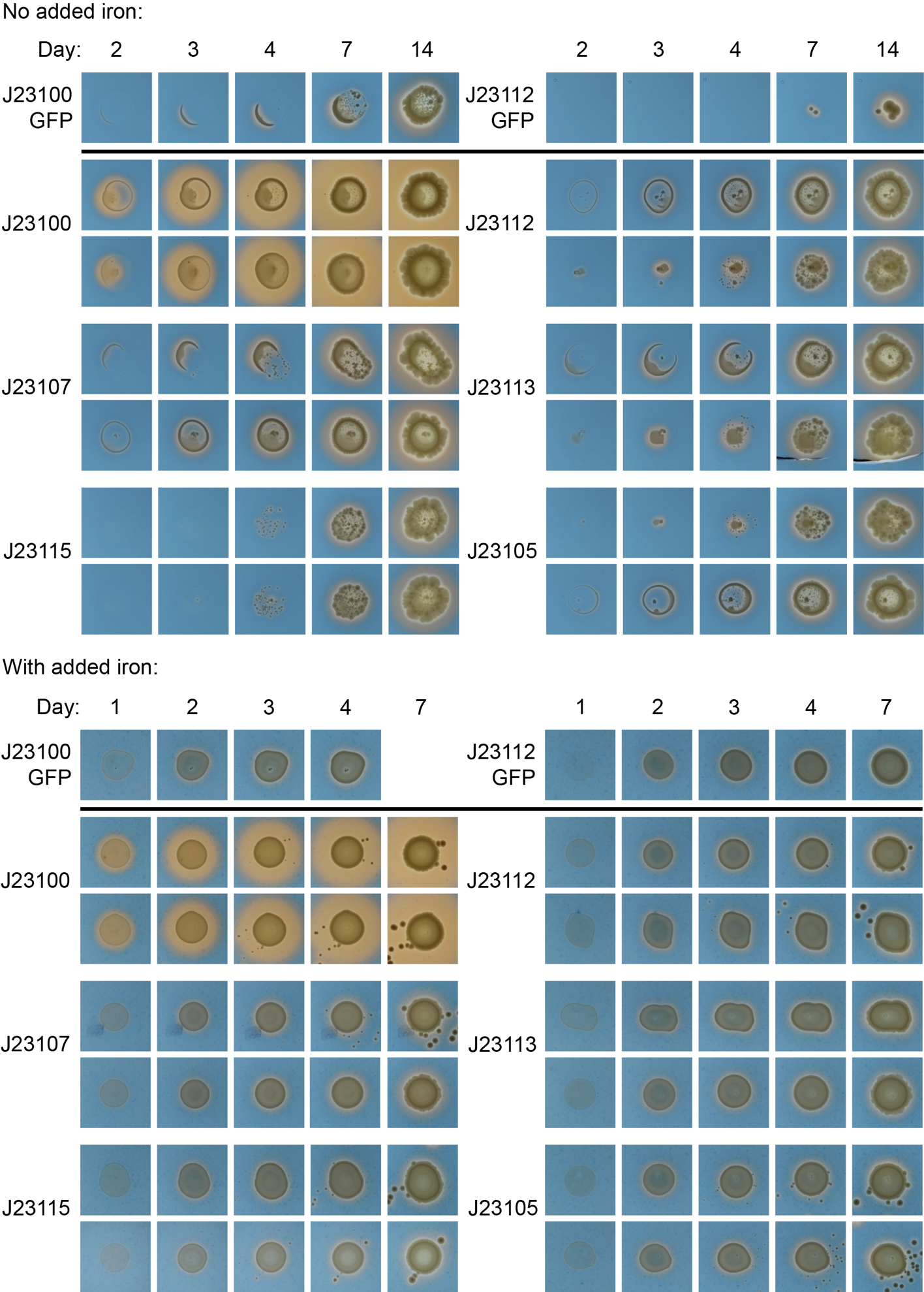

**Fig. S7. CAS agar assay of strains carrying plasmids.** Control cells were transformed with plasmids that express GFP using a synthetic promoter (above the line). Test cells were transformed with plasmids that express the *asb* operon using a synthetic promoter (below the line). Cells were grown on CAS agar plates and imaged intermittently for 7-14 days. Yellow signal indicates sequestration of iron from the blue dye by secreted siderophores.

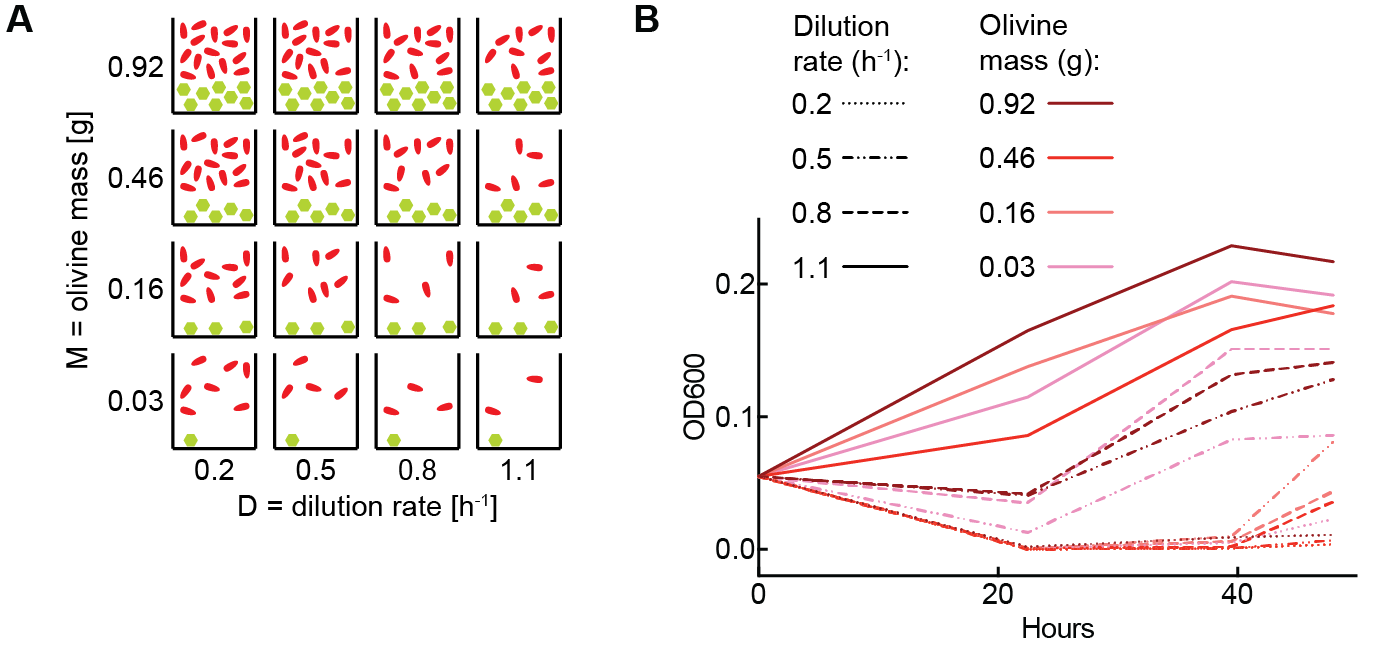

**Fig. S8. Mineral chemostats with the asb+P strain.** A) Experimental layout of parallel chemostats. This strain was tested in the same process conditions as the WT and ∆asbAB strains. B) Optical density of cultures over time after inoculation.

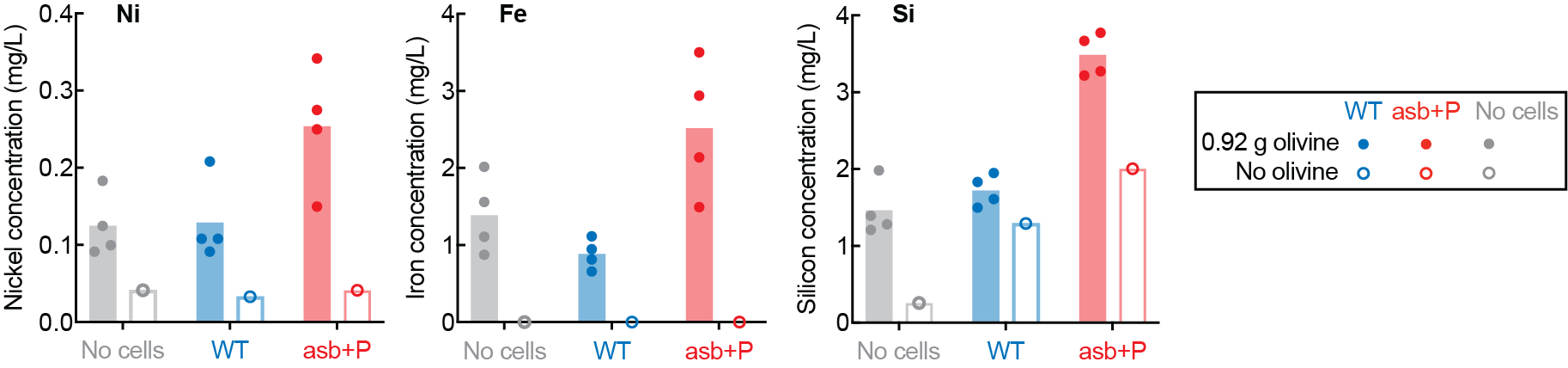

**Fig. S9. Comparison of olivine dissolution rates.** Steady state total concentration of three metals in the culture fluid, measured by ICP-OES.

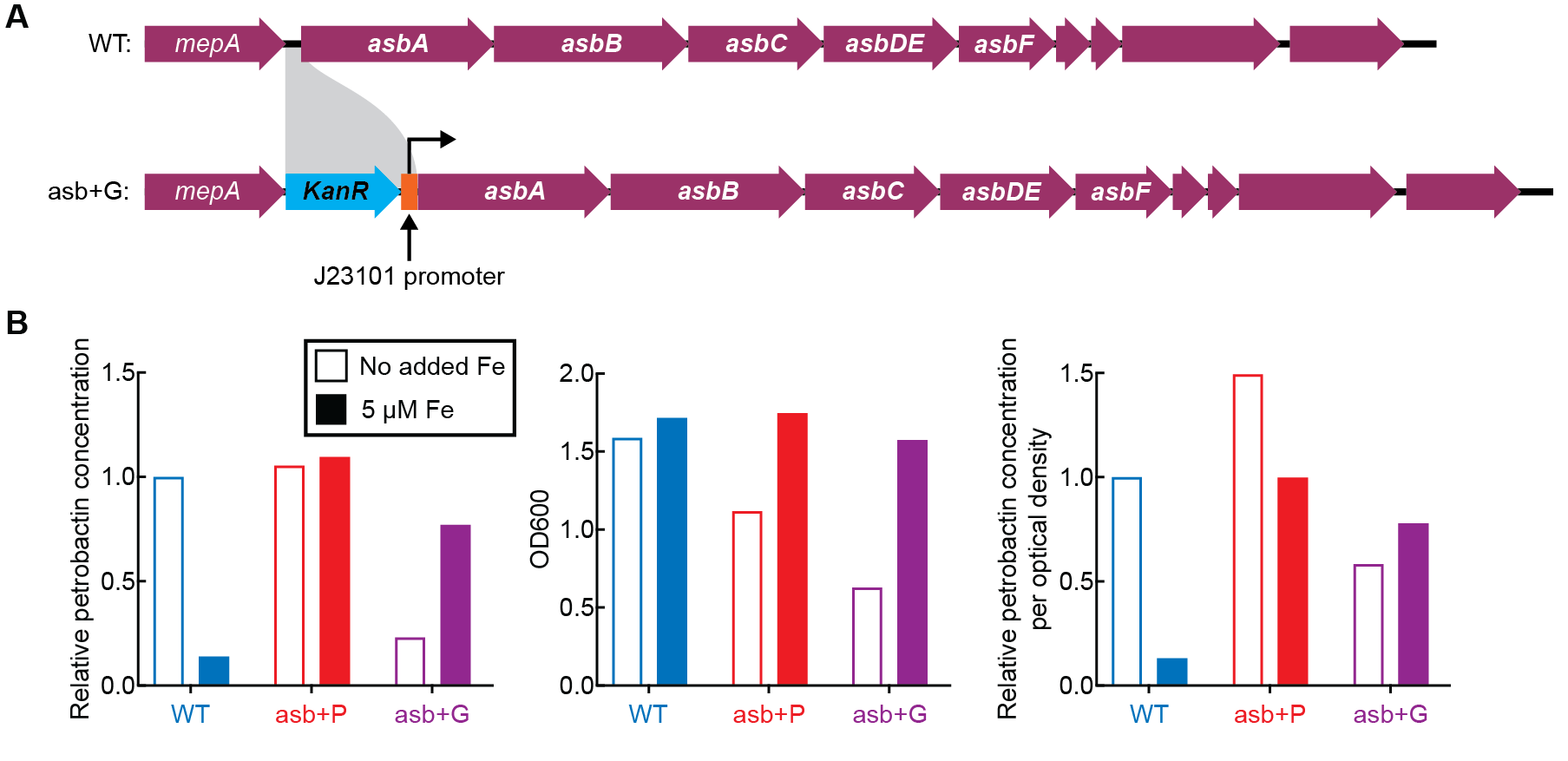

**Fig. S10. Engineering of *A. macleodii* for stable siderophore expression.** A) Construction of the asb+G strain. The native promoter for the *asb* operon was replaced with the synthetic J23101 promoter downstream of a kanamycin resistance marker. B) Production of the siderophore petrobactin in batch shake flasks with or without iron for 24 hours. Cells were inoculated with optical density ~0.01. Graphs show the petrobactin concentration measured by LCMS and normalized to the WT flask with no iron (left), the stationary phase optical density of each flask (middle), the concentration of petrobactin relative to the optical density normalized to the WT flask with no iron (right).

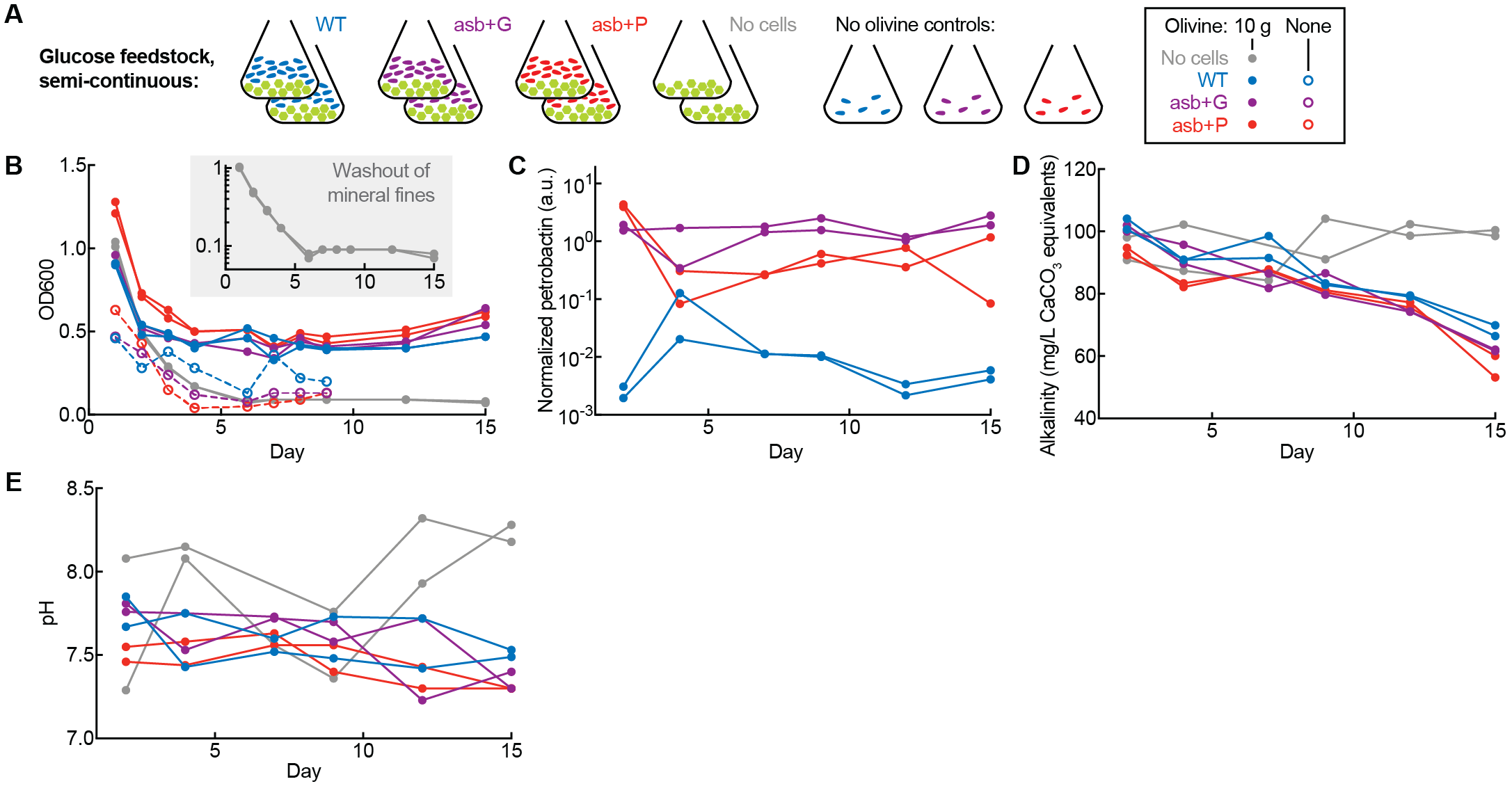

**Fig. S11. Flask-scale weathering with glucose feed.** A) Experimental layout of continuous flask cultures. Every day, 100 mL of the 150 mL culture volume was removed and replaced with fresh media. Flasks with cells contained synthetic medium without buffer. Flasks with no cells contained only synthetic seawater. Flasks with the asb+P strain contained kanamycin. Control flasks with no olivine were stopped after nine days. B) Optical density of flasks over time after inoculation. Flasks with no cells (grey) are shown alone in the inset to visualize the washout of mineral fines. C) Concentration of the siderophore petrobactin, measured by LCMS and normalized within the experiment. D) Alkalinity of filtered culture supernatants over time, measured by automatic titration. E) pH of filtered culture supernatants over time.

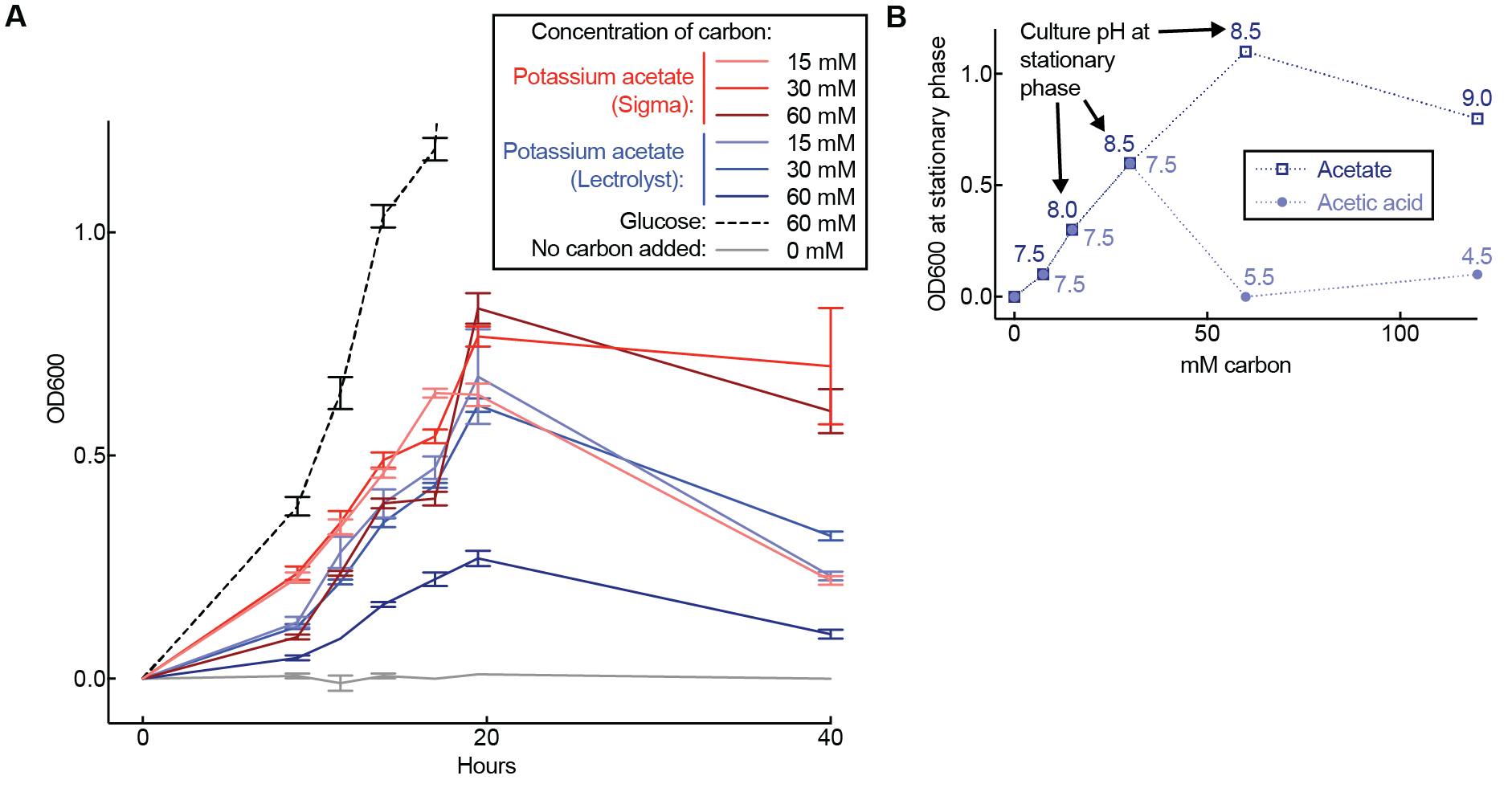

**Fig. S12. Batch growth of *A. macleodii* on acetate**. A) Growth of 4 mL plate cultures measured by optical density at 600 nm. Cells were grown with buffered seawater medium (pH 7.4 at the start of the culture) with different sources of potassium acetate. B) Separate plate cultures to assess the difference between growth acetate and growth on acetic acid. Stationary phase was defined as the maximum cell density measured during the experiment. pH was measured at the end of the cultivation using pH strips. Cultures with high concentrations of acetic acid exhibited no growth because the buffer was immediately overwhelmed.

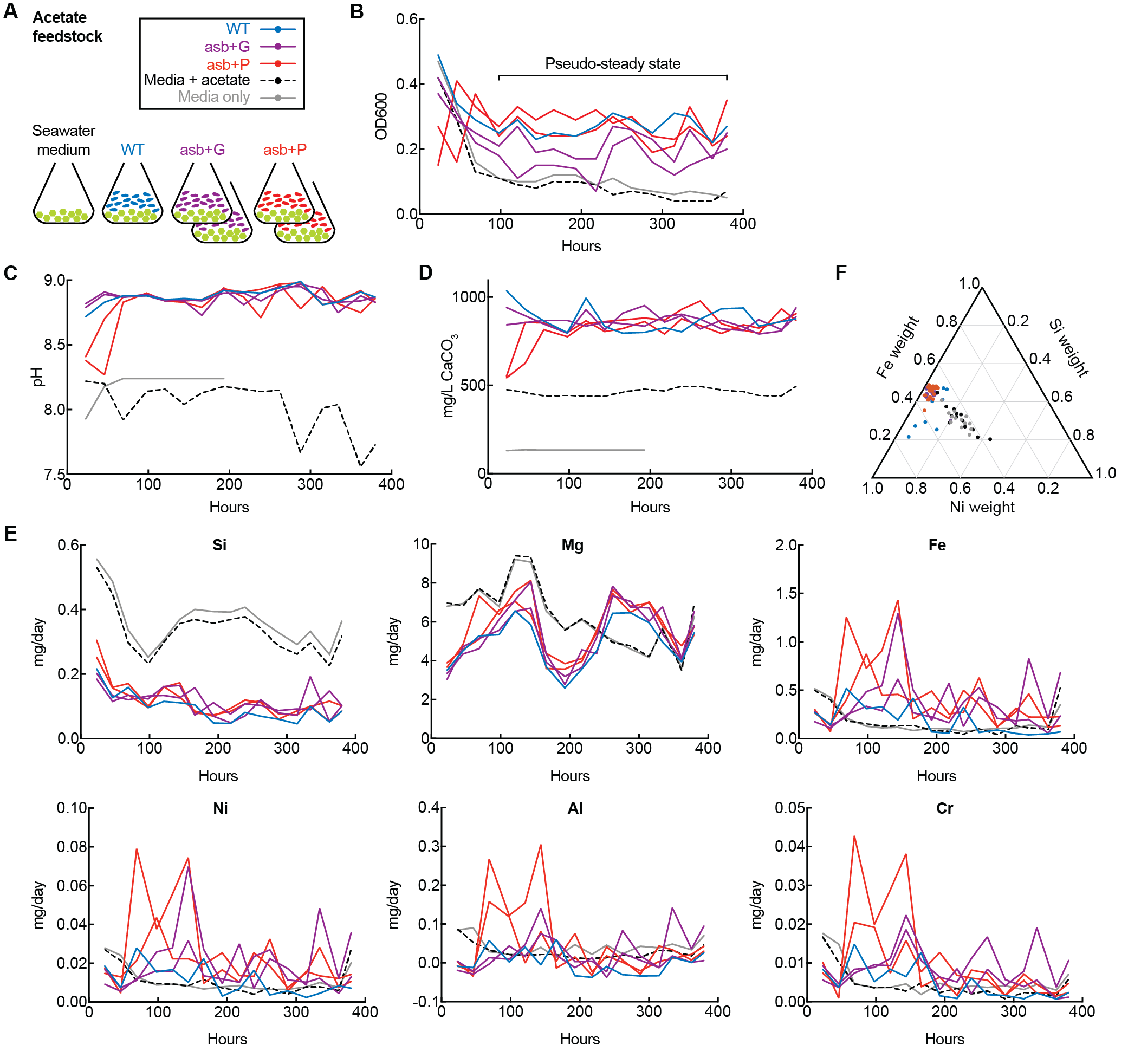

**Fig. S13.** **Flask-scale weathering with acetate feed.** A) Experimental layout of flask reactors fed with acetate. Every day, 100 mL of the 150 mL culture volume was removed and replaced with fresh media. B) Cell density over time measured by OD600. Cultures reached a pseudo-steady state after ~4 days. C) pH of filtered supernatants. D) Total alkalinity of filtered supernatants measured by automatic titration. E) Rate of release of metals from the olivine substrate over time, measured by ICP-OES. F) Relative stoichiometry of mineral dissolution. The concentration of each metal was normalized by the known mass fraction of the metal in the olivine substrate. In the ternary plot, only Ni, Fe, and Si contribute to weighting.

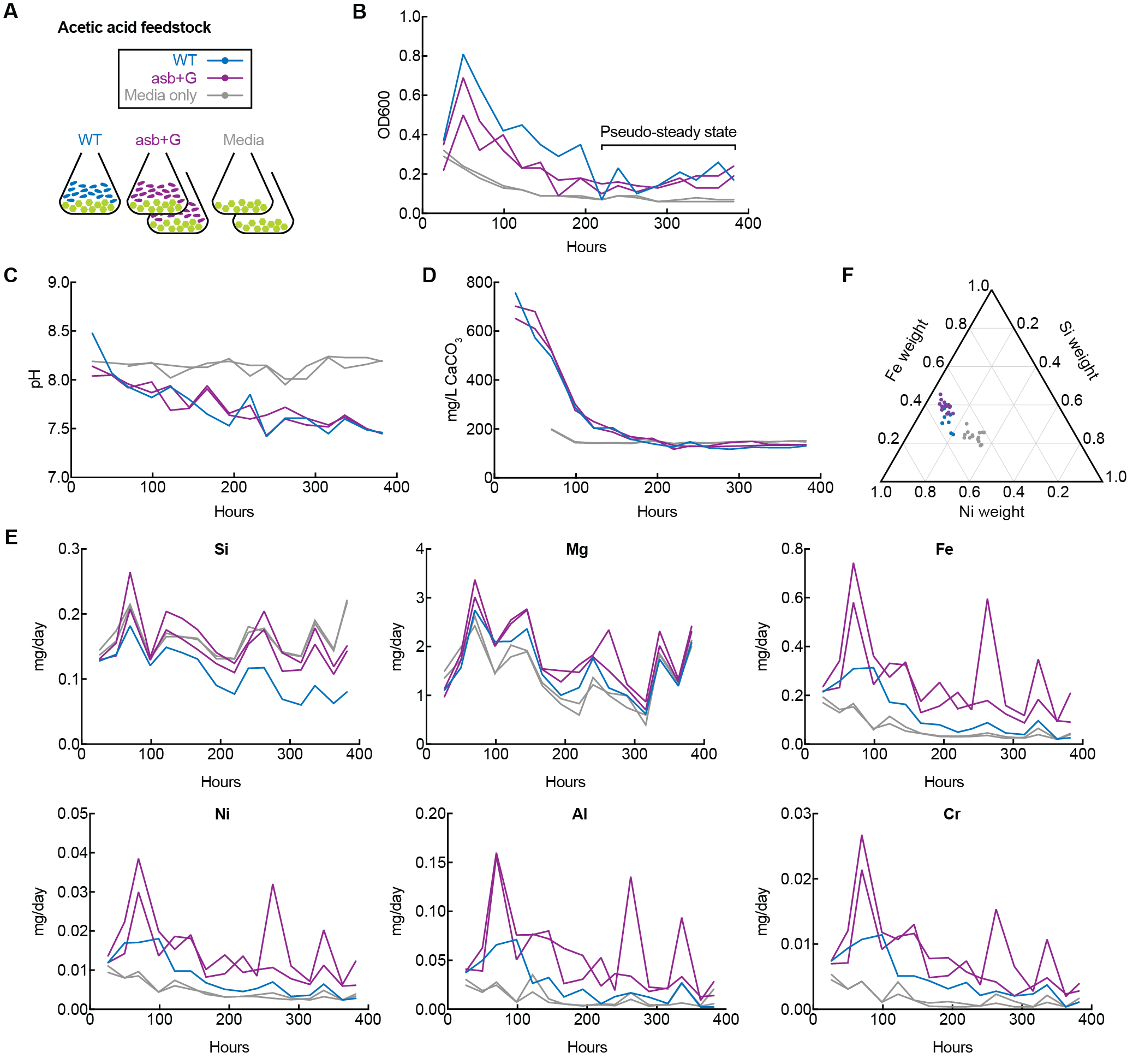

**Fig. S14. Flask-scale weathering with acetic acid feed.** A) Experimental layout of flask reactors. To avoid pH swings, cultures were initiated with acetate. Then, acetic acid media was continuously replaced at 50 mL/day. B) Cell density over time measured by OD600. Cultures reached a pseudo-steady state after ~9 days. C) pH of filtered supernatants. D) Total alkalinity of filtered supernatants measured by automatic titration. E) Rate of release of metals from the olivine substrate over time, measured by ICP-OES. F) Relative stoichiometry of mineral dissolution. The concentration of each metal was normalized by the known mass fraction of the metal in the olivine substrate. In the ternary plot, only Ni, Fe, and Si contribute to weighting.

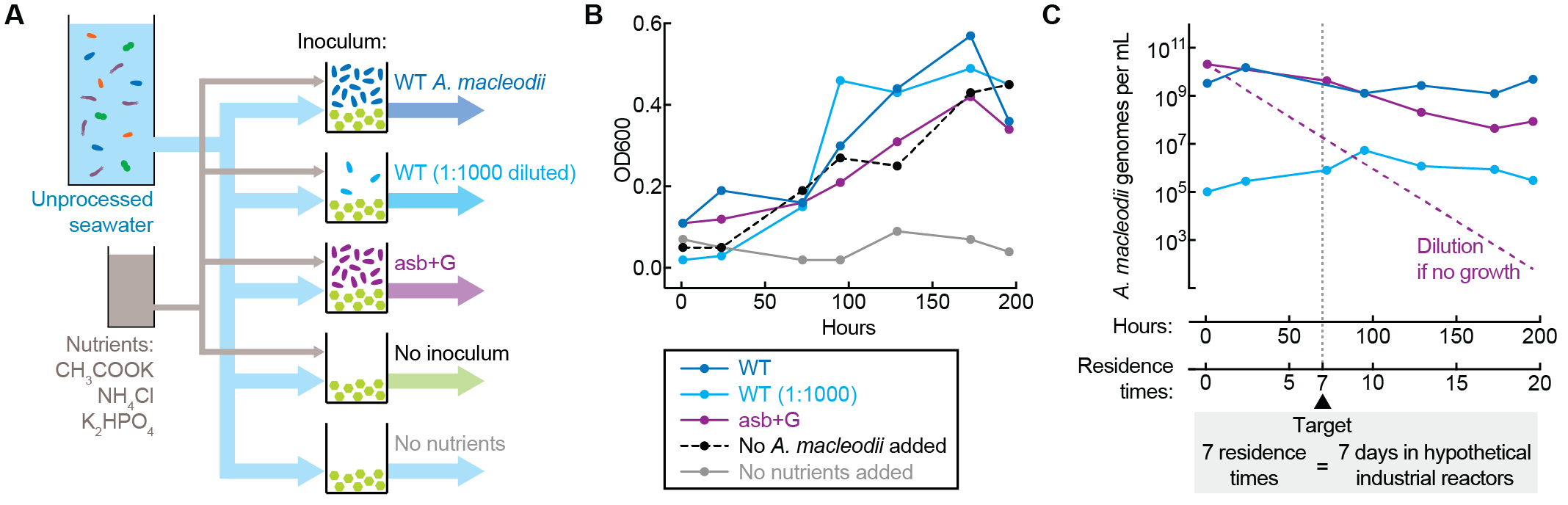

**Fig. S15. Stability of *A. macleodii* in unprocessed seawater.** A) Operation of small-scale mineral reactors with unprocessed seawater. Seawater was flowed through each reactor at rate of 1.98 mL/h. Nutrient solution (14.9 g/L potassium acetate, 3.8 g/L ammonium chloride, 0.6 g/L dipotassium phosphate) was flowed through each reactor at a rate of 0.22 mL/h. With both flows, the residence time of each reactor was 10 h. Vials were inoculated with a 1:20 dilution of saturated *A. macleodii* batch cell culture (OD600 ~ 1) or a dilution thereof. B) Cell density in continuous reactors over time, measured by OD600. C) Abundance of *A. macleodii* DNA in cultures over time, measured by qPCR.

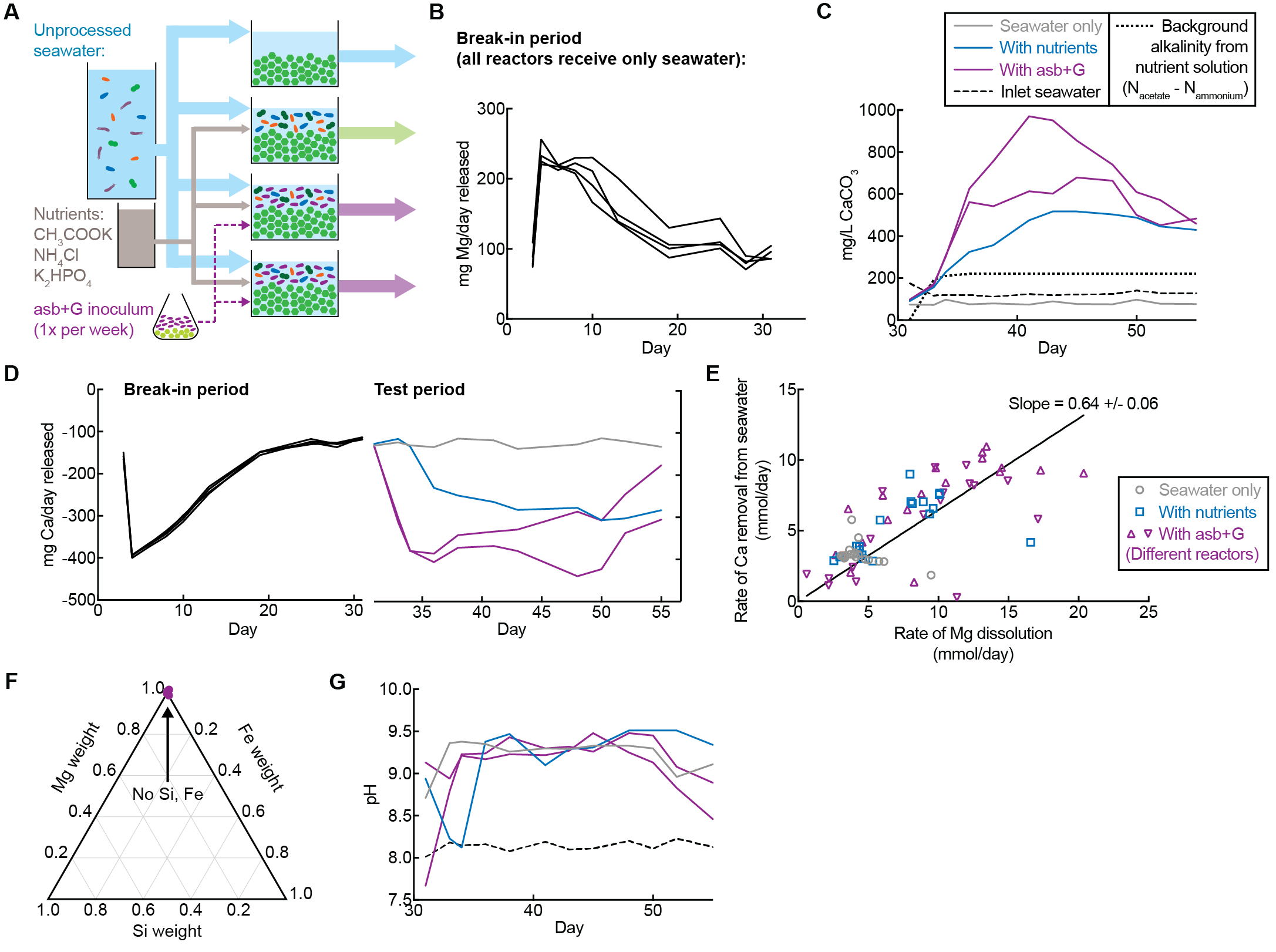

**Fig. S16. Pilot-scale bio-weathering.** A) Experimental layout of pilot mineral bioreactors. Reactors were operated with only seawater for one month, then nutrients were added. B) Rate of release of soluble Mg from the olivine substrate during the one-month break-in period, measured by ICP-OES. C) Total alkalinity of filtered reactor fluid and the inlet seawater over time, measured by automatic titration. Background alkalinity of the nutrient feed was calculated from the proton-accepting capacity of acetate and ammonium in the nutrient solution. D) Rate of release of soluble Ca during the break-in period and of total Ca during the test period, measured by ICP-OES. Negative release rate indicates that the metal was in net removed from the seawater medium and remained in the reactor. E) Plot of Ca and Mg release on a molar basis during the test period. F) Relative stoichiometry of mineral dissolution. The concentration of each metal was normalized by the known mass fraction of the metal in the olivine substrate. In the ternary plot, only Mg, Si, and Fe contribute to weighting. Data not visible is hidden behind visible data points. G) pH of filtered reactor fluid and inlet seawater during the test period.

Appendix 1: Model of mineral chemostats

**Model description**

We modeled the continuous growth of cells in mineral bioreactors using chemostat equations. In a traditional chemostat, cell growth depends on the concentration of a limiting nutrient [X] in the liquid medium. The growth rate is determined by the flow rate of the medium through vessel:

$$\frac{dN}{dt}=\mu_{m}\left( \frac{X}{K_{X}+X} \right)N-DN$$

$$\frac{dX}{dt}=DX_{0}-{\frac{1}{Y_{NX}}\mu}_{m}\left( \frac{X}{K_{X}+X} \right)N-DX$$

$$N=cell density \left[ mass \right]$$

$$\mu_{m}=maximum growth rate \left[ \frac{1}{time} \right]$$

$$X=free concentration of limiting nutrient \left[ \frac{mass}{volume} \right]$$

$$X_{0}=concentration of limiting nutrient in inlet media [X]$$

$$K_{X}=Monod growth constant \left[ X \right]$$

$$D=dilution rate \left[ \frac{1}{time} \right]$$

$$Y_{NX}=yield \left[ \frac{mass of cells}{mass of limiting nutrient} \right]$$

Traditionally, the limiting nutrient in a chemostat is a macronutrient such as glucose.^12^ Chemostats have also been designed to limit growth by the availability of a micronutrient such as iron.^13^ In our model, cell growth depends on both the concentration of glucose [G] and iron [F]. Glucose enters in the liquid medium; in our experiments, we selected G_0_ such that the yield of cells grown with excess iron would be limited by the concentration of glucose in the liquid medium (Fig. S2A). Iron enters the system by dissolution from the olivine substrate at a constant rate. Thus, cell growth at steady state can be limited primarily by one nutrient or the other depending on the chemostat design and operation.

$$\begin{aligned} \frac{dN}{dt}=\mu_{m}\left( \frac{G}{K_{G}+G} \right)\left( \frac{F}{K_{F}+F} \right)N-DN\#\left( 1 \right) \end{aligned}$$

$$\begin{aligned} \frac{dG}{dt}=DG_{0}-{\frac{1}{Y_{NG}}\mu}_{m}\left( \frac{G}{K_{G}+G} \right)\left( \frac{F}{K_{F}+F} \right)N-DG\#\left( 2 \right) \end{aligned}$$

$$\begin{aligned} \frac{dF}{dt}=MR_{F}-{\frac{1}{Y_{NF}}\mu}_{m}\left( \frac{G}{K_{G}+G} \right)\left( \frac{F}{K_{F}+F} \right)N-DF\#\left( 3 \right) \end{aligned}$$

$$M=solid substrate \left[ mass \right]$$

$$R_{F}=solid dissolution rate \left[ \frac{mass of iron}{\left( mass of solid substrate \right)*time} \right]$$

In the context of this study, we are primarily interested in cellular production of siderophores (P), which is known to depend on the extracellular concentration of iron. Torres et al., for example, modeled the yield of siderophores per cell mass as:

$$Y_{PN}=Y_{PNmax}\left( \frac{K_{P}}{K_{P}+F} \right)$$

$$K_{P}=concentration for half maximal production \left[ \frac{iron mass}{volume} \right]$$

$$Y_{PNmax}=Maximum yield\left[ \frac{mass of siderophores}{mass of cells} \right]$$

Which we incorporate into our model as:

$$\begin{aligned} \frac{dP}{dt}=Y_{PNmax}\left( \frac{K_{P}}{K_{P}+F} \right)\mu_{m}\left( \frac{G}{K_{G}+G} \right)\left( \frac{F}{K_{F}+F} \right)N-DP\#\left( 4 \right) \end{aligned}$$

**Parameter selection**

We chose parameters to model the small scale chemostats describe in Fig. 2.

The concentration of glucose in the feed was 5 mM.

- G_0_ = 0.9 g/L

The mineral dissolution rate was approximated from Fig. S9.

- R_F_ = 3 x 10^-6^ g iron/g olivine/h

Maximal growth rate was selected based on our experience with *A. macleodii* cultures and commonly used values for heterotrophic bacteria.

- µ_m­_ = 1.4 h^-1^

The Monod constants for iron and glucose limitation are chosen from published experimental measurements in bacteria. The Monod constant for glucose limited growth for *E. coli* ranges from 50 ug/L to 5 mg/L.^14^ In the case of iron, the Monod constants are more variable and range from 0.1ug/L to 10ug/L. Given this high variation, we chose Monod constants based on the goodness of fit to our data.

- *K_G_* *=* 1x10^-2^ g glucose/L
- *K_F_ =* 1x10^-6^ g iron/L

The yield of biomass from glucose consumption was calculated from experiments in Fig. 2. Specifically, we focused on chemostats with M = 0.92 g olivine. With low dilutions rates (D = 0.2, 0.5) we did not observed expression of *asbA*, suggesting that cells do not experience iron limitation:

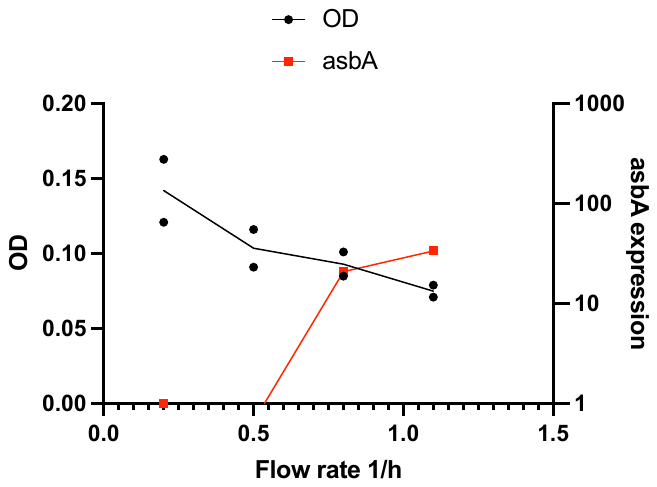

We assumed that glucose is nearly completely consumed (G ≈ 0) when D = 0.2. At steady state, equations (1) and (2) combine to:

$$\begin{aligned} 0=G_{0}-\left( \frac{1}{Y_{NG}} \right)N \end{aligned}$$

We calculated the yield using by assuming that an optical density of 1 (OD600 = 1) confers 0.56 g biomass /L.

- Y_NG_ = 0.09

The yield of biomass from iron was calculated similarly from experiments with M = 0.032 g olivine, for which the expression of *asbA* was always high, suggesting that cells experience iron limitation:

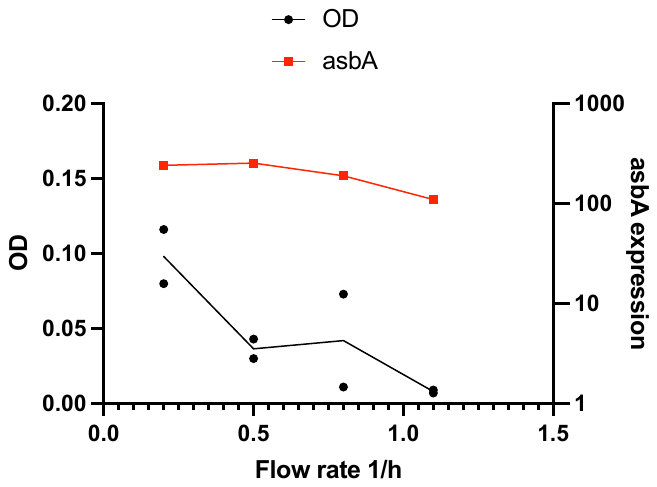

We assumed that iron is nearly completely consumed (F ≈ 0) when D = 0.2. At steady state, equations (1) and (3) combine to:

$$\begin{aligned} 0=MR_{F}-\left( \frac{1}{Y_{NF}} \right)DN \end{aligned}$$

We calculated the yield using by assuming that an optical density of 1 (OD600 = 1) confers 0.56 g biomass /L.

- Y_NF_ = 7 x 10^4^

The iron concentration of half maximal siderophore repression K_p_ was set equivalent to K_F_, consistent with our understanding of siderophore production only under iron limitation. The siderophore yield Y_PNmax_ was fit to experimental data.

To determine which nutrient was primarily limiting (Fig. 2B), we calculated the following value:

$$\begin{aligned} \left( \frac{G}{K_{G}+G} \right)^{-1}\left( \frac{F}{K_{F}+F} \right) \end{aligned}$$
